# Lipid nanoparticle encapsulated nucleoside-modified mRNA vaccines elicit polyfunctional HIV-1 antibodies comparable to proteins in nonhuman primates

**DOI:** 10.1101/2020.12.30.424745

**Authors:** Kevin O. Saunders, Norbert Pardi, Robert Parks, Sampa Santra, Zekun Mu, Laura Sutherland, Richard Scearce, Maggie Barr, Amanda Eaton, Giovanna Hernandez, Derrick Goodman, Michael J. Hogan, Istvan Tombacz, David N. Gordon, R. Wes Rountree, Yunfei Wang, Mark G. Lewis, Theodore C. Pierson, Chris Barbosa, Ying Tam, Xiaoying Shen, Guido Ferrari, Georgia D. Tomaras, David C. Montefiori, Drew Weissman, Barton F. Haynes

## Abstract

Development of an effective AIDS vaccine remains a challenge. Nucleoside-modified mRNAs formulated in lipid nanoparticles (mRNA-LNP) have proved to be a potent mode of immunization against infectious diseases in preclinical studies, and are being tested for SARS-CoV-2 in humans. A critical question is how mRNA-LNP vaccine immunogenicity compares to that of traditional adjuvanted protein vaccines in primates. Here, we found that mRNA-LNP immunization compared to protein immunization elicited either the same or superior magnitude and breadth of HIV-1 Env-specific polyfunctional antibodies. Immunization with mRNA-LNP encoding Zika premembrane and envelope (prM-E) or HIV-1 Env gp160 induced durable neutralizing antibodies for at least 41 weeks. Doses of mRNA-LNP as low as 5 μg were immunogenic in macaques. Thus, mRNA-LNP can be used to rapidly generate single or multi-component vaccines, such as sequential vaccines needed to protect against HIV-1 infection. Such vaccines would be as or more immunogenic than adjuvanted recombinant protein vaccines in primates.

## Introduction

Messenger ribonucleic acid (mRNA)-based vaccines have been shown to elicit protective immunity against Zika virus (ZIKV) infection after a single immunization of rhesus macaques ^1^, and have been designed for many other pathogens including Ebola, influenza, Hepatitis C, Cytomegalovirus, and respiratory syncytial virus ^2–7^.

Recent development of mRNA vaccines has overcome initial roadblocks to their use. mRNA vaccines were initially hindered by innate immune sensing of mRNA ^7–9^. However, advances in modifying mRNA nucleosides has improved mRNA translation, while eliminating recognition by innate pattern recognition receptors ^7,8,10^. Specifically, the incorporation of pseudouridine and 1-methyl-pseudouridine prevents recognition of mRNA by Toll-like receptor 7 and 8 and other nucleic acid sensing pattern recognition receptors ^7^. Modification of mRNA in combination with novel HPLC and FPLC purification methods and codon optimization methods have increased the efficiency of mRNA production and the efficacy of mRNA vaccination ^8,11,12^. For mRNA to be effective at transducing cells in vivo, it must be protected from RNAses. One method for protecting the mRNA from degradation has been its encapsulation in lipid nanoparticles (LNP) ^7,8,13^. Importantly, small interfering RNAs (siRNAs) in LNP have been approved by the FDA for treatment of a genetic form of amyloidosis ^14^. Thus, advances in encapsulation and preventing mRNA immune sensing have made mRNA vaccines a feasible vaccine platform.

mRNA vaccines are also attractive as a vaccine platform, because they can accept the genes of various pathogen antigens, and they can be rapidly manufactured at scale. Theses two aspects made mRNA vaccines a prime candidate for responding to the SARS-CoV-2 outbreak ^15,16^. High levels of SARS-CoV-2 neutralizing antibodies, defined as higher than geometric mean values of convalescent serum from COVID-19 patients, were observed for both nucleoside-modified mRNA-LNP and Matrix-M1 adjuvanted subunit protein in human trials^16–18^. Given the ability of mRNA vaccines to elicit neutralizing antibodies ^8,16,17^, this platform could improve the elicititation of neutralizing antibodies against HIV-1. However, little data exists on the comparative immunogenicity of mRNA-LNP and proteins in non-human primates. Thus, a key question for HIV-1 vaccine development is whether the immunogenicity of mRNA vaccines encoding HIV-1 immunogens, a poorly immunogenic protein that requires extensive post-translational modifications, is comparable to that of proteins that can be purified after in vitro production.

HIV-1 vaccines aiming to elicit protective antibody responses will likely need to elicit polyfunctional non-neutralizing effector antibodies (nnAbs) and/or broadly neutralizing antibodies (bnAbs) ^19,20^. NnAbs are easy to induce and bind to the surface of virus-infected CD4+ T cells where they can mediate antibody-dependent cellular cytotoxicity (ADCC). However, their efficacy in protecting against HIV transmission or control of the disease in the setting of high transmission rates was called into question with the HVTN 702 ALVAC-C, gp120-C trial failing to show efficacy ^21,22^. The only HIV-1 clinical trial to date to demonstrate any efficacy, RV144, induced nnAbs by immunizing with gp120 proteins derived from HIV-1 isolates A244 and MN ^23^. Protection in that trial correlated with binding IgG to the second variable region on Env and ADCC, but the protective antibody response was not durable ^24^.

In contrast to nnAbs, bnAbs are difficult to induce for many reasons including shielding of bnAb Env epitopes by glycans, the metastability of Env conformation, conformational masking of neutralizing epitopes ^25–27^, host immune controls preventing bnAb development ^28,29^, and the requirement for multiple highly improbable somatic mutations that are needed for acquisition of antibody neutralization breadth ^30–32^. mRNA immunization elicits antigen-specific follicular T helper cells, which are key for affinity maturation of antibodies in germinal centers ^33,34^, and has been shown to correlate with bnAb development during natural infection ^35,36^. Regardless of whether it is nnAbs or bnAbs targeted, the poor durability of HIV-1 Env antibody responses^37^, and the necessity of designing a vaccine that has multiple components for bnAb induction^19,20^ necessitates novel vaccine platforms that can maintain protective antibody levels.

Here, we compare the antibody responses in rhesus macaques induced by either HIV-1 Env-encoded as nucleoside-modified mRNA-LNP or the same vaccine candidate administered as an adjuvanted HIV-1 Env recombinant protein. In each comparison, mRNA-LNP administration induced equal or better immune responses as proteins. Moreover, mRNA-LNP vaccination encoding soluble HIV-1 Env trimers or ZIKV prM-E elicited durable neutralizing antibodies that were stable for approximately one year.

## Results

### Both protein and mRNA vaccination elicited high titers of serum HIV-1 binding antibodies

We compared the immunogenicity of HIV-1 Env A244 gp120 lacking the first 11 N-terminal amino acids of gp120 (Δ11 gp120, as in the RV144 trial) ^38^ administered as either nucleoside-modified mRNA-LNP or adjuvanted recombinant protein in rhesus macaques. We immunized two groups of five macaques with recombinant protein formulated in different adjuvants. One group of five macaques received A244 Δ11 gp120 formulated with the aluminum hydroxide adjuvant Rehydragel (**Fig. 1A**). This group mimicked the A244 Δ11 gp120 protein aluminum hydroxide (Rehydragel) formulation used in the RV144 ALVAC/B/E gp120 trial ^23^. Adjuvants vary in their immunostimulatory strength; thus, in a second group of five macaques, we assessed the immunogenicity of the A244 Δ11 gp120 Env in combination with a liposomal adjuvant formulation composed of monophosphoryl lipid A (a TLR-4 agonist) and QS21 (ALFQ) ^39^. Additionally, two groups of macaques were immunized with nucleoside-modified mRNA-LNP encoding different forms of A244 Δ11 gp120. One group of macaques received A244 Δ11 gp120, whereas the other group received mRNAs encoding A244 Δ11 gp120 with a D368R mutation that disrupted the CD4 binding site (**Fig. 1A**). The purpose of the CD4 binding site knockout mutation was to determine whether Env binding to CD4^+^ T cells *in vivo* altered immunogenicity of HIV-1 Env gp120. Macaques were immunized with either protein or mRNA-LNP at weeks 0 and 6 and antibody responses were followed for 18 total weeks (**Fig. 1A**).

**Figure 1.**
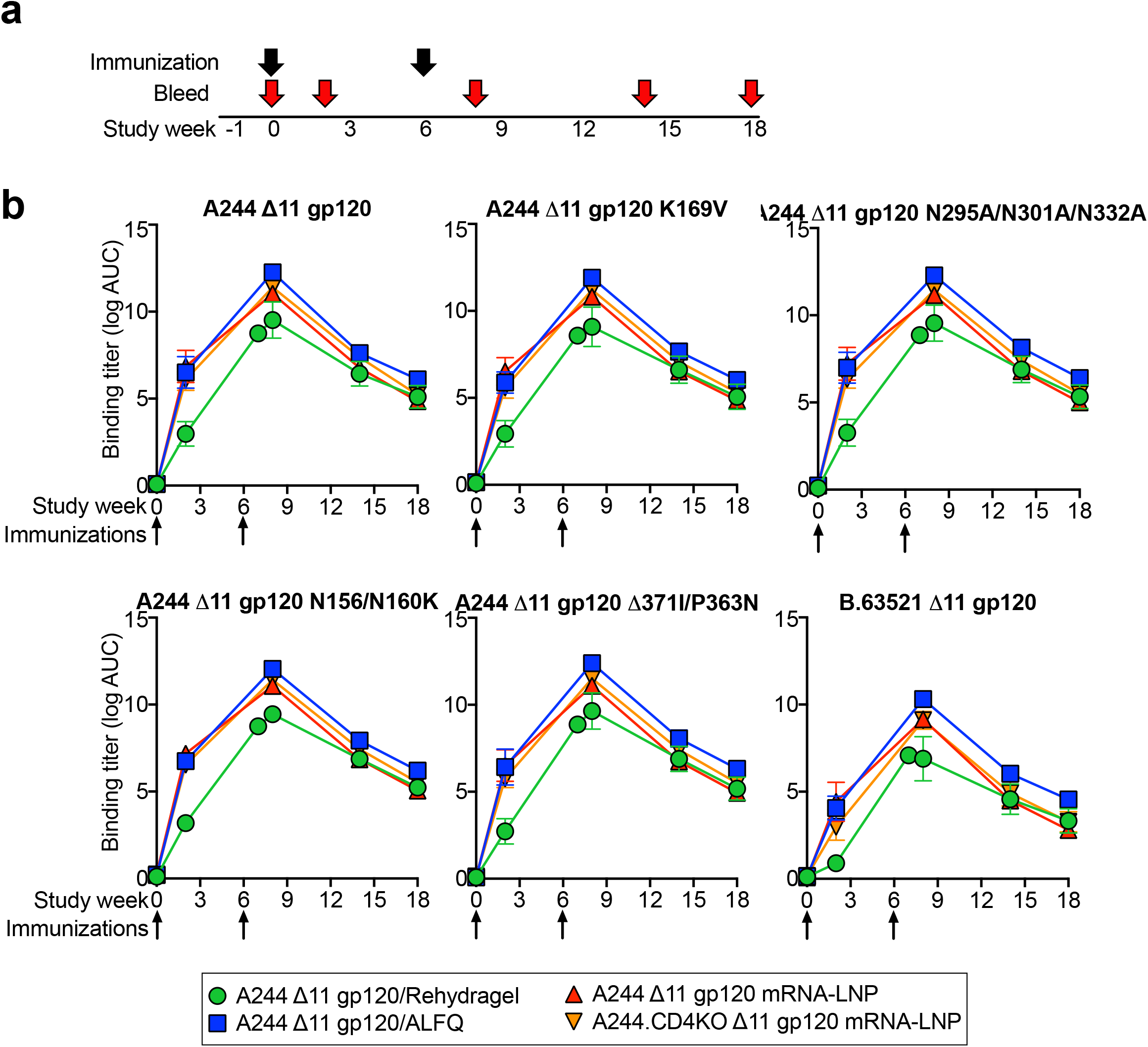
Immunization of rhesus macaques with HIV-1 Env recombinant protein adjuvanted with ALFQ or HIV-1 Env-encoding mRNA-LNP elicits comparable titers of gp120-specific serum IgG. **a**. Rhesus macaque vaccine regimen and biospecimen collection schedule. The different immunogens are listed on the right side. **b**. Serum IgG binding titers for each group are shown as mean log area-under-the-curve (AUC) against A244 gp120, A244 gp120 mutants, and B.63521 gp120. A244 gp120 was mutated at the ADCC site of immune pressure observed in the RV144 trial (A244 K169V). Additionally, the V3-glycan (N295A/N301A/N332A), V2-glycan (N156K/N160K), CD4 binding site (delta371I/P363N) neutralizing epitopes were mutated on A244 gp120. The group mean and standard error are shown (n = 5 per group). Arrows indicate immunization timepoints.

Binding nnAbs to Env immunogens have been associated with protection in macaques from SHIV infection ^40,41^. Thus, we first determined serum binding IgG elicited by each of the mRNA/LNP or protein groups. Regardless of site of immunization, gp120-specific IgG titers in the macaque serum were detectable after a single immunization and continued to rise after second immunization (**Fig. 1B**). mRNA-LNP elicited nearly identical serum IgG antibody titers as compared to macaques immunized with A244 Δ11 gp120 formulated with ALFQ (**Fig. 1B**). Comparison of mRNA-LNP that encoded wildtype gp120 versus CD4 mutated gp120 showed identical binding IgG elicitation (**Fig. 1B**). Macaques that received A244 Δ11 gp120 formulated with Rehydragel generated the lowest group mean serum IgG binding titers to A244 Δ11 gp120. For these ELISA assays, the differences between the Rehydragel group and the other groups were greatest after one immunization, but the small numbers of monkeys per group precluded statistical analysis. Nonetheless, ALFQ adjuvant induced higher levels of A244 Δ11 gp120 antibody compared to gp120 in Rehydragel (**Fig. 1B**). The same group ranking was also observed for binding IgG to a HIV-1 Env gp120 B.63521 from a non-vaccine matched HIV-1 isolate (**Fig. 1B**). The binding of serum antibodies to A244 gp120 was not reduced by the introduction of mutations in the CD4 binding, glycan-V3, and glycan-V2 sites (**Fig. 1B**). Thus, mRNA-LNP gp120 vaccination produced comparable IgG titers in plasma as recombinant gp120 protein in the potent adjuvant ALFQ.

IgG responses to the second variable (V2) region of HIV-1 Env were a correlate of reduced infection risk in the RV144 trial ^24^. Similarly, K169 in the V2 region was a site of immune pressure during the RV144 trial ^42^. To assess whether nucleoside-modified mRNA-LNP vaccination elicited comparable V2 region IgG antibodies in plasma, we tested serum IgG binding to various minimal antigens that recapitulated the V2 region of HIV-1 Env. These antigens included V1V2 proteins, a V2 peptide, and V2 scaffolded on gp70. Binding to each antigen showed the same pattern as gp120 binding. Recombinant protein adjuvanted with AFLQ and mRNA-LNP immunization elicited nearly identical plasma IgG responses (**Fig. 2**). Recombinant gp120 adjuvanted with Rehydragel elicited the weakest responses for all the antigens tested, with group difference being largest after a single immunization (**Fig. 2**). V1V2-specific binding IgG titers were comparable whether the V1V2 antigen matched the vaccine strain or if it was from an unmatched virus 1086C (**Fig. 2A**). The first variable region (V1) was not necessary for V2 binding, as plasma IgG bound to a short V2 only peptide, K178 (**Fig. 2B**). The antigen for which binding IgG correlated with reduced infection risk in the RV144 trial was B.CaseA V1V2 scaffolded on gp70 ^24,43^. Each group of macaques generated binding IgG antibodies to gp70 B.CaseA V1V2 protein (**Fig. 2C**). The antibodies were specific for HIV-1 V1V2, as no antibodies were detected against the control gp70 scaffold presenting murine leukemia virus V1V2 (**Fig. 2C**). However, mutation of K169 had little effect on B.CaseA V1V2 binding IgG titers. Similarly, mutation of the K169 in the V2 region of A244 gp120 had little effect on binding IgG titers (**Fig. 1B**). Thus, V1V2 antibodies were not solely dependent on the K169 at the site of immune pressure observed in the RV144 trial.

**Figure 2.**
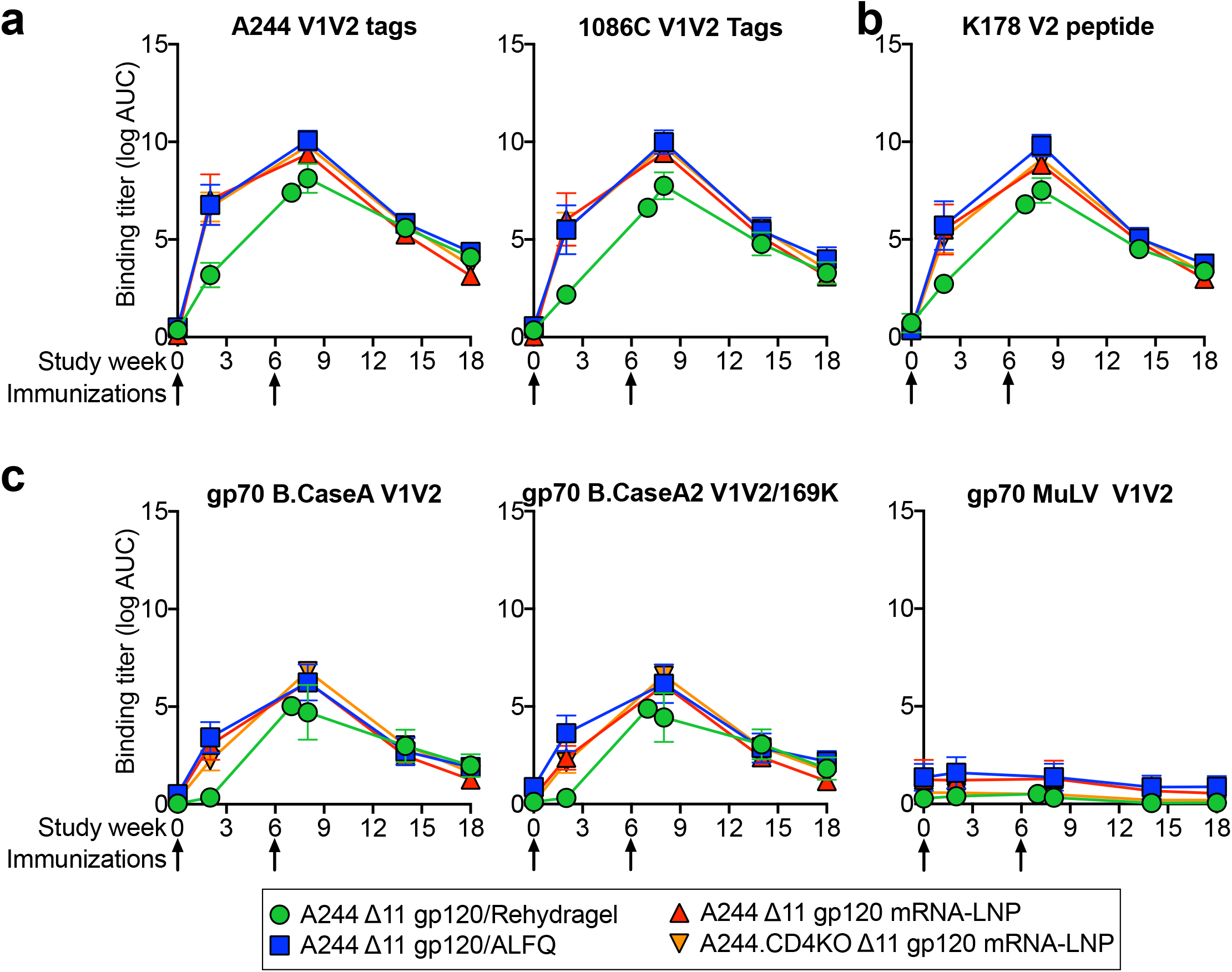
Adjuvanted recombinant protein and mRNA-LNP immunization of rhesus macaques elicits gp120 V2-specific plasma IgG responses. Plasma IgG binding titers recombinant V1V2 proteins, V2 peptide and gp70 scaffolded V1V2 proteins. The group mean and standard error are shown (n = 5 per group). Arrows indicate immunization timepoints.

To determine the breadth of reactivity and epitope specificity of the first and second variable (V1V2) region antibodies induced by each vaccine, we performed linear V1V2 peptide arrays. Vaccination induced only low levels of V1-binding antibodies, but high titers of V2 antibodies across the different vaccines (**Fig. 3A-D and Supplementary Fig. 1**). V2 antibody binding magnitudes were similar for mRNA-LNP and protein formulated with ALFQ (**Fig. 3**). Each vaccine elicited antibodies capable of binding V2 peptides from clade AE and C (**Fig. 3 and Supplementary Fig. 1**). Therefore, mRNA-LNP and adjuvanted protein vaccination elicited V2 antibody responses that were similar in magnitude, breadth, and epitope specificity.

**Figure 3.**
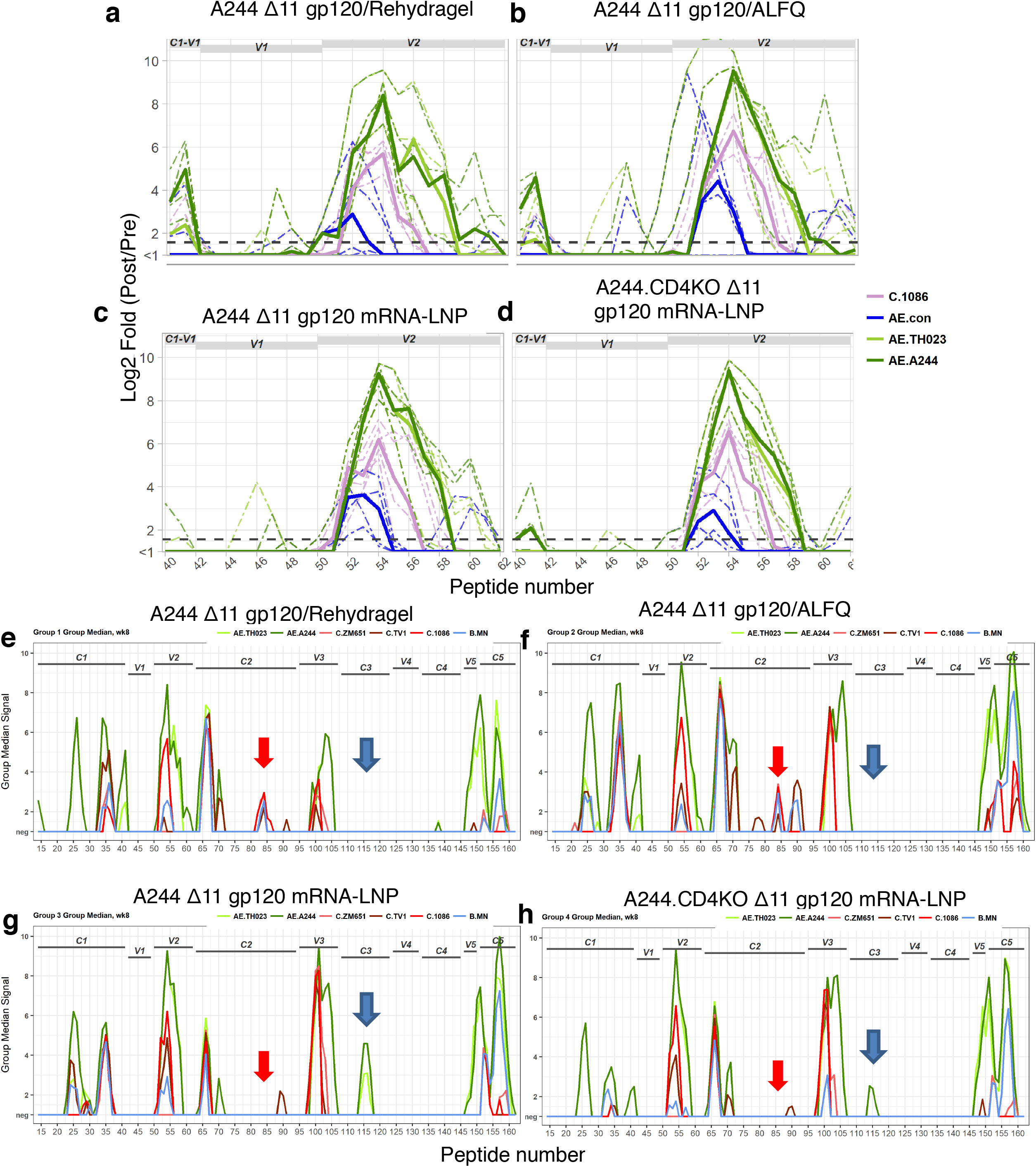
Adjuvanted recombinant gp120 and mRNA-LNP-encoded gp120 immunizations elicit plasma IgG with similar V2 specificities, but different C2 and C3 specificities. Plasma IgG binding to HIV-1 Env peptides spanning (**a-d**) the V1V2 region of gp120 or (**e-f**) the entire gp120 subunit. Arrows indicate differences in C2 (red) and C3 (blue) binding antibodies. HIV-1 Env regions encompassed by the peptides are listed above the curves. Each panel shows the response for the immunization group listed in the title. Graph lines are colored based on HIV-1 isolate. The group medians are shown by solid lines. Dashed lines indicate individual macaques in **a-d**.

Next, we compared plasma antibody specificities induced by protein or mRNA-LNP immunization at HIV-1 Env gp120 sites outside of the V1V2 site. mRNA-LNP induced similar A244 gp120 linear epitope antibodies compared to protein formulated in adjuvant with two exceptions. First, recombinant protein, but not mRNA-LNP, elicited plasma IgG to the C-terminal portion of the second constant (C2) region (**Fig. 3E-H and Supplementary Fig. 2**). Second, mRNA-LNP immunization elicited plasma IgG against the third constant region but recombinant protein immunization did not (**Fig. 3E-H and Supplementary Fig. 2**).

To further compare polyclonal plasma IgG specificities, we assessed the ability of post-vaccination plasma to block the binding of gp120 monoclonal antibodies (mAbs) to HIV-1 Env. Plasma from either adjuvanted protein- or mRNA-LNP-immunized macaques was added to HIV-1 Env protein, followed by the addition of biotinylated mAbs to determine plasma antibody blocking of monoclonal antibody binding to Env. We examined blocking of non-neutralizing effector antibodies CH58, which targets the V2 site of immune pressure identified in RV144 ^44^, and A32 which defines an immunodominant ADCC gp120 site ^45^ that synergizes with V2 antibodies to mediate ADCC ^46^ (**Fig. 4A**). Next, we determined whether plasma could block the HIV-1 entry receptor CD4 from binding to Env (**Fig. 4B**), or block V2-glycan bnAbs PG9 and CH01 and N332 glycan-dependent bnAbs 2G12 and PGT125 Env binding (**Fig. 4C,D**). Either immunization with adjuvanted recombinant gp120 protein or mRNA-LNP elicited plasma antibodies that blocked the binding of CH58, A32, CD4, CH01, 2G12, and PGT125 (**Fig. 4**). Comparison of mRNA-LNP-induced antibodies versus gp120 protein-induced plasma antibodies showed that mRNA-LNP immunization elicited the same magnitude of blocking after 2 immunizations as protein adjuvanted with ALFQ (**Fig. 4**). Blocking activity was lowest for animals immunized with protein adjuvanted with Rehydragel (**Fig. 4**). The blocking of CH01 binding to A244 most likely represented CH58-like antibody binding and not V2-glycan bnAb binding, since CH01 binding can be blocked by CH58-like linear V2 antibodies (**Fig. 4**).

**Figure 4.**
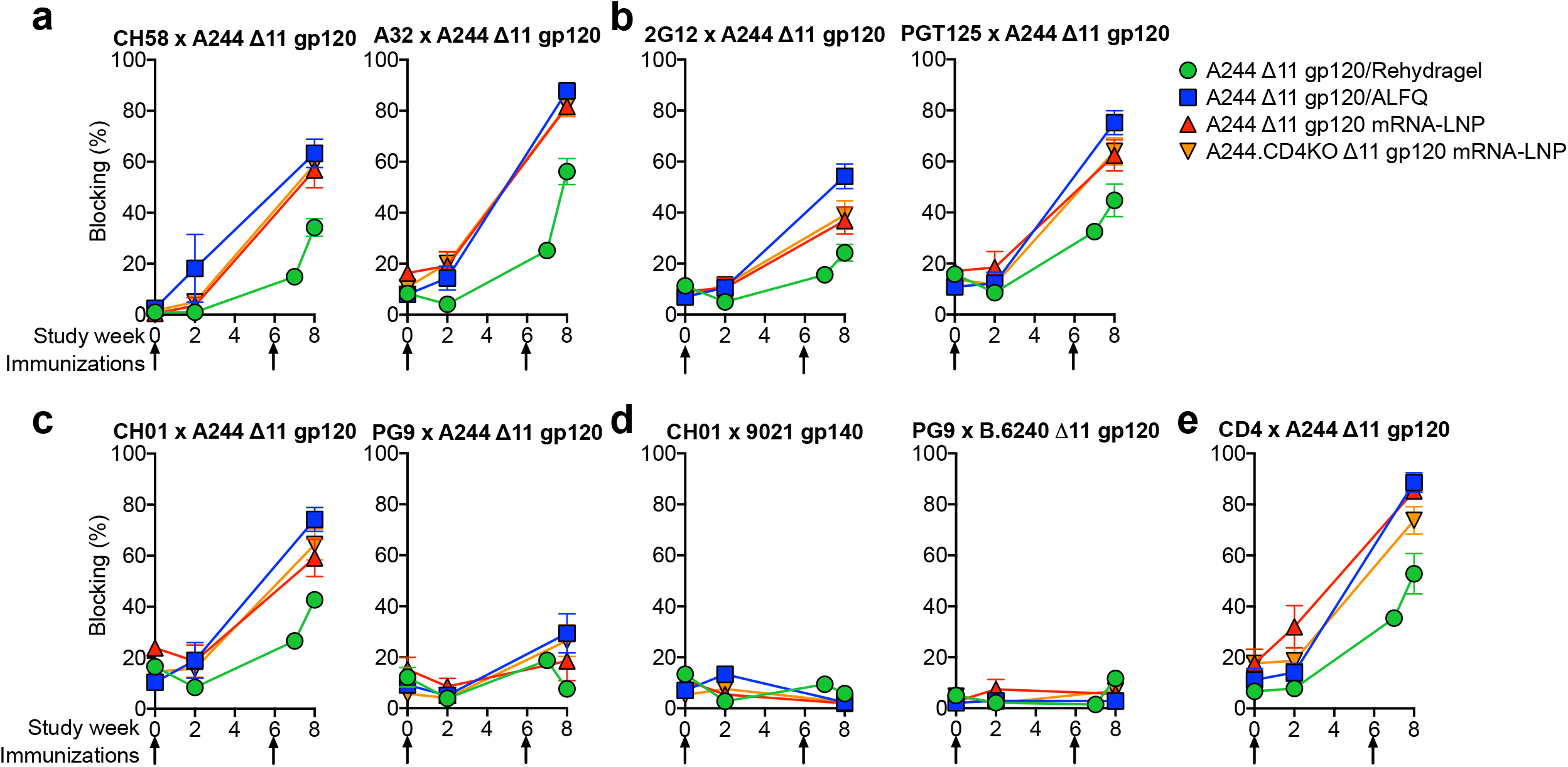
ALFQ-adjuvanted recombinant gp120 and mRNA-LNP-encoded gp120 immunizations in macaques elicit similar magnitudes of plasma antibodies capable of blocking ADCC-mediating antibodies, HIV-1 bnAbs, and CD4 binding to Env. **a-e.** Plasma antibody blocking of (**a**) ADCC-mediating antibodies (CH58 and A32), (**b**) N332 glycan bnAbs (2G12 and PGT125), (**c,d**) V2-glycan CH01, PG9), and (**e**) soluble CD4. Antibody and Env names are listed in the graph title. Note, plasma IgG blocking is absent when non-vaccine matched Envs 9021 or B.6240 are used as the Env antigen in **d**. The group mean and standard error are shown (n = 5 per group). Arrows indicate immunization timepoints.

### Comparable neutralizing and non-neutralizing antibody functions induced by adjuvanted Env gp120 protein versus mRNA-LNP immunization

Several IgG Fc receptor (R)-mediated immune responses have been associated with decreased transmission risk either in the RV144 trial or in Env immunizations studies in macaques followed by low dose mucosal SHIV challenges ^41,47^. These immune responses include binding to HIV-1 infected CD4^+^ T cells, ADCC and antibody-dependent cellular phagocytosis (ADCP).To assess the potential for week 8 plasma IgG to mediate effector functions against infected cells, we first examined plasma IgG binding to HIV-1.CM235-infected T cells. Plasma IgG from all four groups of macaques was able to bind to HIV-infected cells (**Fig. 5A,B**). When binding levels of IgG was quantified as mean fluorescence intensity of bound IgG or the percentage of cells positive for HIV-1 protein p24 and plasma IgG, mRNA-LNP vaccination and recombinant protein adjuvanted with ALFQ were not different (**Fig. 5A,B**). In agreement with the overall lower IgG titers, Rehydragel-adjuvanted protein gave the weakest cell binding responses (**Fig. 5A,B**).

**Figure 5.**
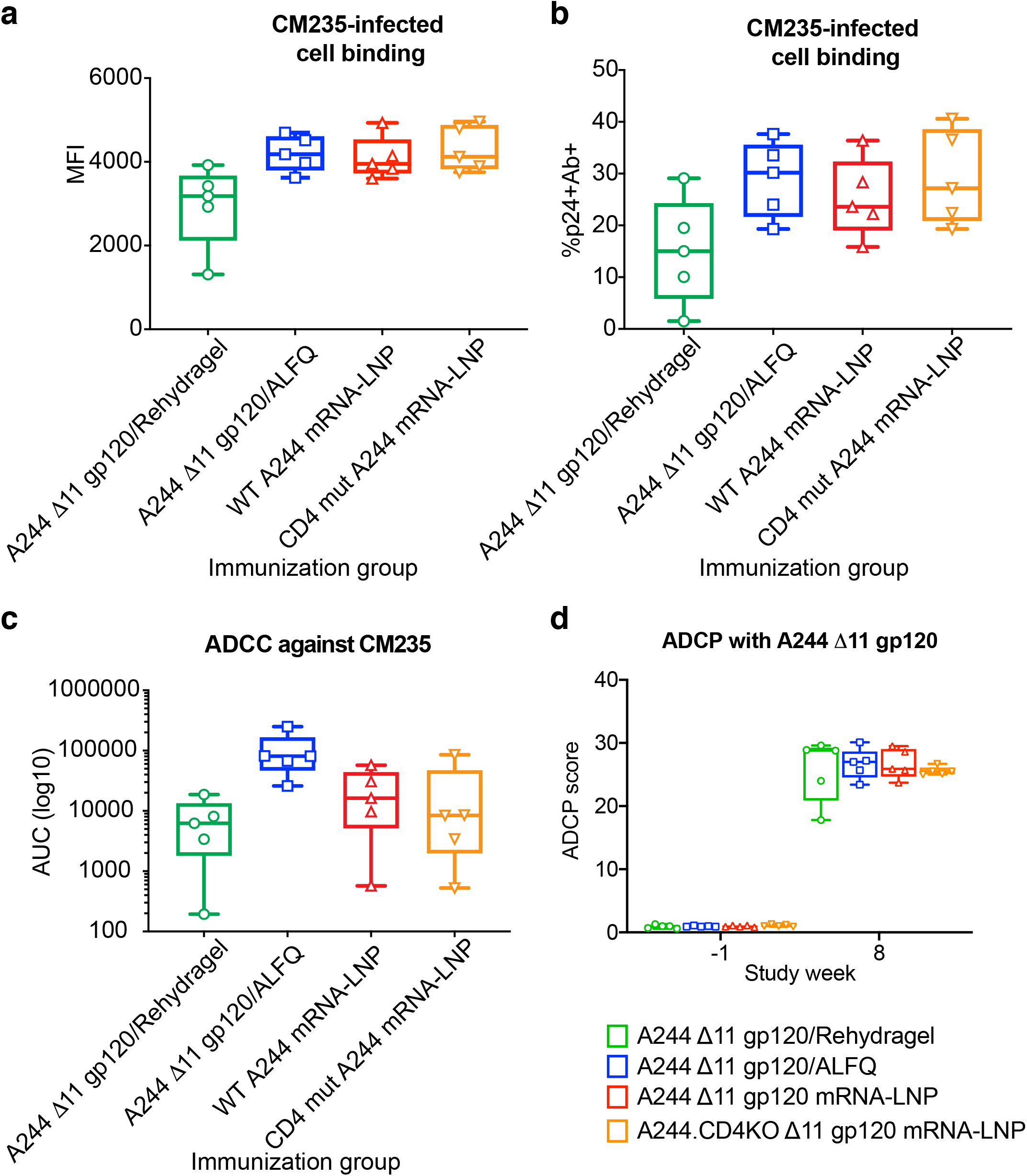
Adjuvanted recombinant protein and mRNA-LNP immunizations elicit antibodies that mediate comparable magnitudes of infected cell binding and effector functions. **a,b**. Plasma IgG binding to CM235-infected cells measured as mean fluorescence intensity (MFI) or percentage of infected (p24+) cells with detectable plasma IgG binding (Ab+). Each symbol represents an individual macaque. **c.** Antibody-dependent cellular cytotoxicity (ADCC) of CM235-infected cells. **d**. Antibody-dependent cellular phagocytosis (ADCP) of A244 Δ11 gp120-coated fluorescent beads. Symbols indicate scores for individual macaques immunized with adjuvanted recombinant A244 Δ11 gp120 protein (green and blue) or A244 Δ11 gp120 mRNA-LNP (red and orange). HIV-1 bnAb 2G12 and influenza bnAb CH65 were used as positive and negative controls respectively. Box and whisker plots show minimum values, maximum values, median, and interquartile ranges.

ADCC activity was measured in a flow cytometry-based granzyme B assay. There was a clear adjuvant effect between the two protein immunized group. ADCC titers were approximately one order of magnitude higher when ALFQ was used as the adjuvant as compared to Rehydragel. Similarly, mRNA-LNP immunization (mean±standard deviation = 22,916±22,102) induced activity lower than ALFQ-formulated recombinant protein, but 3-fold higher than protein adjuvanted with Rehydragel (mean±standard deviation = 7,324±7,024) (**Fig. 5C**). Elimination of CD4 binding to A244 Δ11 gp120 had no effect on ADCC activity (**Fig. 5C**). Lastly, we compared ADCP activity of week 8 plasma antibodies from macaques administered adjuvanted recombinant protein or mRNA-LNP. Median plasma ADCP activity against A244 gp120-coated beads was similar across all groups with A244 Δ11 gp120 in Rehydragel eliciting a wider range of responses (**Fig. 5D**). In agreement with previous studies, nucleoside-modified mRNA-LNP vaccination elicited antibody effector functions that have been shown previously to correlate with reduced infection risk ^24,48^.

While the goal of A244 Δ11 gp120 immunization was to induce non-neutralizing effector functions like those seen in the RV144 trial, we compared elicitation of neutralizing antibodies by each of the vaccines. We selected for testing the tier 1 HIV-1 strain CRF_01AE 92TH023, against which A244 has consistently induced neutralizing antibodies ^24,44^. We found that there was no significant difference among the different vaccination regimens for induction of HIV-1 92TH023 neutralizing antibodies (**Supplementary Fig. 3**). These studies showed that for immunization with gp120 monomers aiming to elicit potentially protective non-neutralizing V2 antibodies, nucleoside-modified mRNA-LNP vaccination was superior to recombinant protein formulated with aluminum hydroxide and comparable to recombinant protein formulated with a more complex TLR-4/QS21/liposomal adjuvant.

### Immunizations of macaques with sequential CH505 Env gp160 mRNA-LNP, SOSIP gp140 mRNA-LNP, or adjuvanted recombinant SOSIP gp140 proteins

We previously isolated CD4 binding site bnAbs from the African individual CH505 ^49,50^. From CH505, we cloned a series of different CH505 Envs for testing as immunogens to induce similar types of bnAbs with vaccination ^49,50^. To this end, we studied a mRNA-LNP sequential immunization strategy where each individual envelope in the series is delivered one-at-a-time in a specific order. The HIV-1 envelope immunogens were designed as transmembrane HIV-1 gp160s (50 μg/immunization) or soluble, stabilized gp140 SOSIP trimers (50 μg/immunization; **Fig. 6A**). The SOSIP gp140s were stabilized by a chimeric gp41 and the addition of E64K and A316W mutations ^51^. The immunogenicity of these two sets of mRNA-LNP was compared to that of 100 μg of the same set of CH505 Envs as soluble gp140 SOSIP proteins formulated in the TLR-4 adjuvant, GLA-SE (**Fig. 6A**). Four rhesus macaques were immunized every four weeks with either set of immunogens (**Fig. 6A**), and binding antibody and neutralizing antibodies were measured.

**Figure 6.**
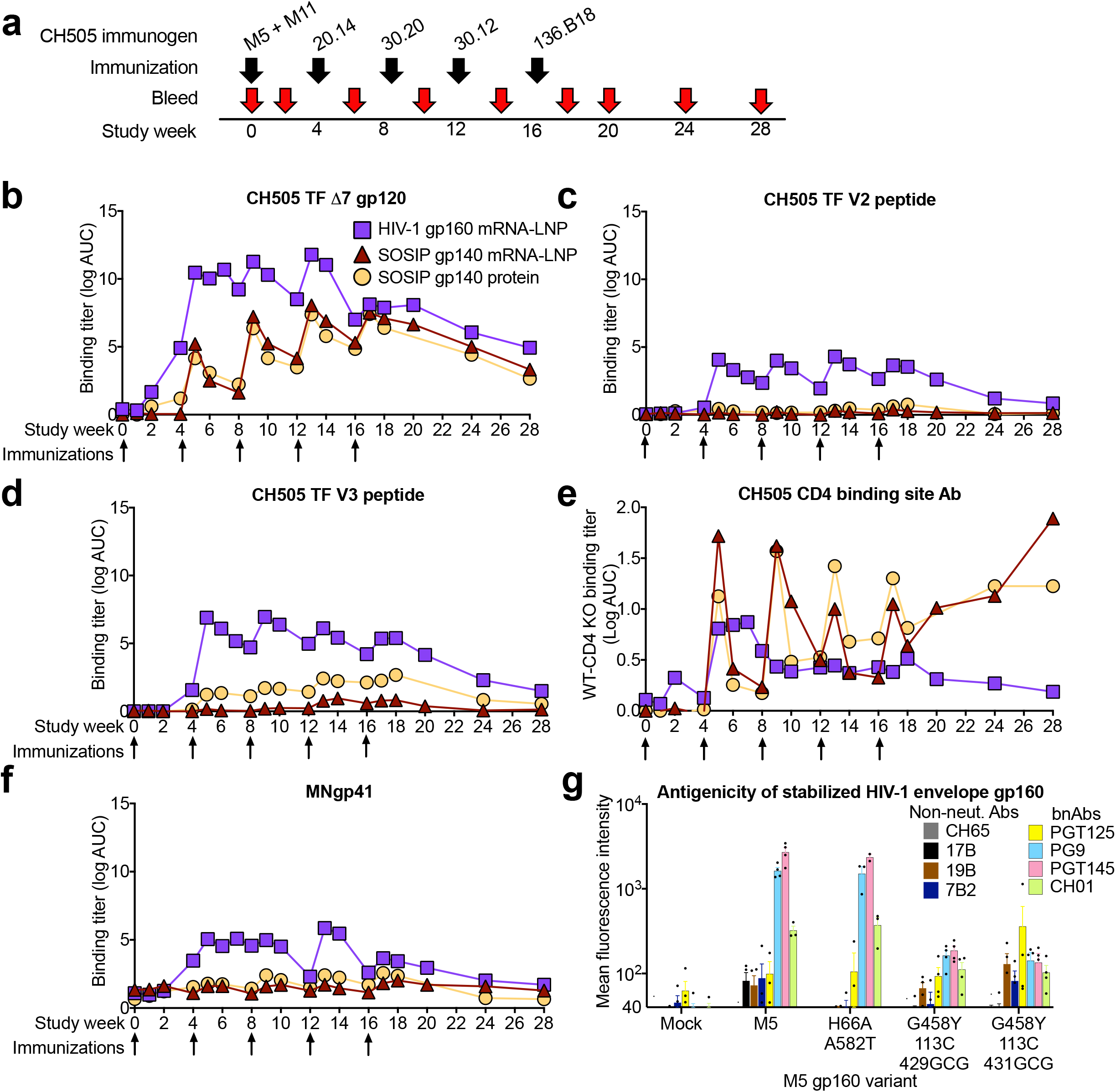
Comparison of HIV-1 Env trimer immunogenicity after mRNA-LNP and recombinant protein immunization. **a.** Macaque immunization regimen and biospecimen collection. **b-d.** Plasma IgG binding titers as log AUC to CH505 TF **b**. gp120, **c**. V2, and **d**. V3. **e**. Difference in binding titers to wildtype CH505 TF gp120 and CH505 TF gp120 with the CD4 binding site knocked out with a Δ371Ile mutation. Values above zero indicate higher binding to wildtype gp120 than the CD4 binding site mutant. **f**. Plasma IgG binding titers as log AUC to HIV-1.MN gp41. Lines represent group mean values (n = 4 per group). **g.** Antibody binding to HIV-1 gp160 expressed on the surface of Freestyle293F cells after mRNA transfection. Antibody binding is shown as mean fluorescence intensity measured by flow cytometry. Dark-colored and bright-colored bars indicate non-neutralizing and broadly neutralizing antibodies respectively. Antibody epitopes examined were: 7B2, gp41; 17B, co-receptor binding site; 19B, V3; PGT145, trimer apex; CH01 and PG9, V1V2-glycan; PGT125, V3-glycan; and CH65, anti-influenza heamaglutttin antibody. Independent replicates are indicated by black circles. Mean and standard error are shown for the 2-4 independent replicates.

All three vaccines were immunogenic in macaques. With regard to induction of binding titers of gp120 antibodies, mRNA-LNP encoding sequential CH505 gp160s induced higher gp120 titers than did either of the mRNA-LNP encoding soluble gp140 SOSIP trimers or soluble gp140 SOSIP trimers proteins (**Fig. 6B**). These binding titers rose dramatically after two immunizations with gp160 mRNA-LNP, but rose gradually over the course of five immunizations in macaques immunized with SOSIP gp140s (**Fig. 6B**). Plasma IgG binding titers to gp120 were equivalent between the mRNA-LNP and the adjuvanted soluble gp140 SOSIP trimer protein-immunized animals (**Fig. 6B**). Soluble gp120 exposes non-neutralizing epitopes, thus we examined whether the high titers of gp120 antibodies in macaques immunized with gp160 mRNA-LNP were due to antibodies targeting these sites. Indeed, the gp160 mRNA-LNP induced very high titers of antibodies to linear CH505 TF V2 peptides, whereas titers in the gp140 SOSIP trimer groups showed no or a slight increase from baseline (**Fig. 6C**). Linear V3 epitopes were also highly immunogenic in gp160 mRNA-LNP-immunized macaques, and to a lesser extent was immunogenic in soluble gp140 SOSIP trimer protein immunized macaques (**Fig. 6D**). Interestingly, V3 antibodies were not elicited by gp140 SOSIP trimer mRNA-LNP, suggesting a difference in V3 exposure when the trimer was expressed *in vivo* or potential effects of the adjuvant. To examine the induction of gp120 antibodies that may target conserved neutralizing epitopes, we assessed the amount of gp120 antibody binding that was dependent on the CD4 binding site. We mutated the CD4 binding site with a deletion of the isoleucine at position 371 and determined the decrease in plasma IgG binding to CH505 TF gp120. Both groups of macaques that received SOSIP trimers as either protein gp140s or mRNA-LNP were superior to the mRNA-LNP encoding the gp160s (**Fig. 6E**). Immunization with gp140 SOSIP trimers elicited differential binding between the mutant and wildtype gp120 that rapidly receded after each immunization, but was maintained for 16 weeks once immunizations were stopped (**Fig. 6E**). Similar to linear V2 and V3 peptide antibodies, gp160 mRNA-LNP induced high levels of gp41 antibodies whereas gp140 SOSIP trimers delivered as mRNA-LNP or adjuvanted recombinant protein had levels close to baseline (**Fig. 6F**). Thus, gp160 mRNA-LNP immunization elicited higher titers of undesired antibodies targeting non-neutralizing gp120 and gp41 epitopes and lower titers of desired CD4 binding site antibodies than SOSIP trimer-encoding mRNA-LNP or adjuvanted recombinant protein. However, none of the regimens induced significant tier 2 or heterologous neutralizing antibodies due to the need for high affinity germline B cell targeting and designs of sequential Env regimens that select for bnAb improbable mutations (**Supplementary Tables 1-3**) ^20,31,32,52,53^.

The high levels of undesired antibodies elicited by gp160 mRNA-LNP immunization could be due to poorly-folded HIV-1 envelope expressed on the surface of cells. We hypothesized that stabilization of HIV-1 envelope with amino acid substitutions that preserve Env trimers in their native conformation could reduce HIV-1 envelope binding to non-neutralizing antibodies. We introduced stabilizing amino acid changes H66A and A582T into envelope ^54^. Also, we generated envelopes with G458Y and cysteines at positions 113 and 429 or 113 and 431 ^55^. The cysteines have been shown to form disulfide bonds that keep the HIV-1 envelope in the closed conformation^55^ and G458Y stabilizes the envelope fifth variable loop^53^. Among these variants of the M5 gp160 envelope, adding H66A and A582T reduced binding by non-neutralizing antibodies against V3 and gp41, while leaving trimer-specific (PGT145) or timer-preferring (PG9 and CH01) antibody binding unchanged (**Fig. 6G**). Thus, mRNA immunization with stabilized HIV-1 envelope gp160 is one potential approach to improve the elicitation of neutralizing antibodies, while not engaging the B cell receptor of B cells that produce antibodies that cannot bind native, fusion-competent HIV-1 envelope.

### Nucleoside-modified mRNA-LNP immunization induces durable antibody responses against Zika prM-E and HIV-1 Env

Inducing durable Env antibody responses is a key goal of HIV vaccine development. Yet in the RV144 trial, protective antibodies fell dramatically over the first 42 weeks after vaccination ^24^. Thus, if immunization with mRNA-LNP induced durable antibody responses it would benefit their use as a vaccine platform for many different infectious diseases. We assessed the durability of antibody responses using neutralization assays of the tier 1 CH505 w4.3 virus, since antibodies capable of neutralization of the tier 2 CH505 transmitted founder (TF) virus were not elicited (**Supplementary Tables 1-3**). CH505 w4.3 is an early virus isolate that is identical to the tier 2 CH505 TF virus with the exception of a W680G mutation that makes it highly sensitive to HIV antibody neutralization ^49,56,57^. Longitudinal comparative analyses of neutralization of the CH505 w4.3 virus showed that the gp160-encoding mRNA-LNP elicited higher titers of neutralizing antibodies than the gp140 SOSIP trimer-encoding mRNA-LNP or the gp140 SOSIP trimer proteins in GLA/SE ^58^ (**Fig. 7A**). Notably, neutralization of the CH505 w4.3 virus was still detectable in all three groups 12 weeks after the last immunization (**Fig. 7A**). There were also sporadic low levels of neutralization of CH505 or 426C viruses that were modified to be highly sensitive to CD4 binding site antibodies (**Supplementary Tables 1-3**).

**Figure 7.**
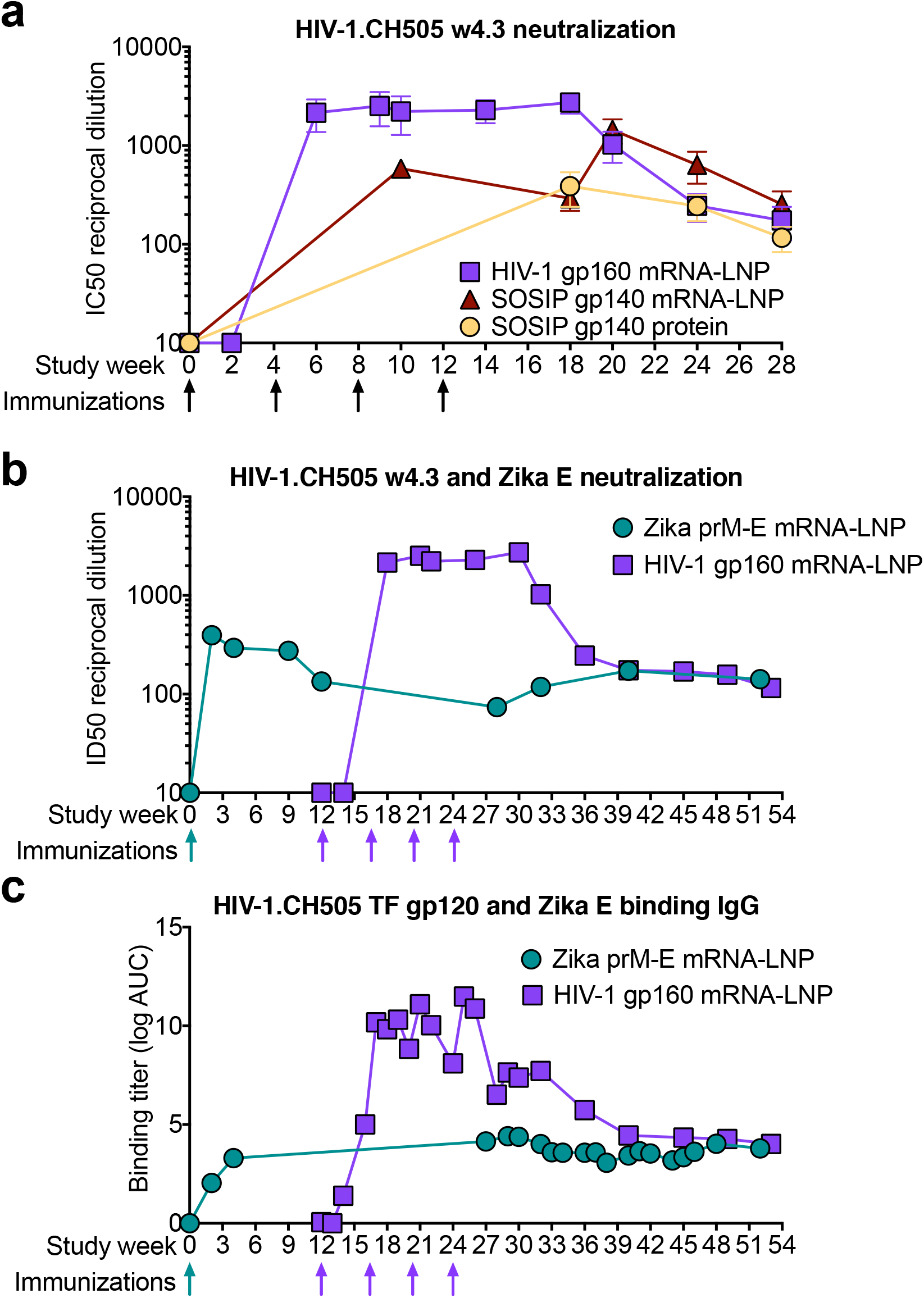
HIV-1 Env mRNA-LNP immunization, like Zika prM-E immunization, elicits durable HIV-1 serum neutralizing antibodies. **a.** Comparison of macaque serum neutralization of HIV-1 infection of TZM-bl cells. Neutralization titers are shown as the reciprocal serum dilution that inhibits 50% of virus replication (ID50). Lines represent the group geometric mean for macaques immunized with gp160 mRNA (purple), SOSIP gp140 mRNA (maroon), or SOSIP gp140 protein (light orange). **b.** Vaccine-elicited serum neutralization titers as ID50 against Zika (forest green) and HIV-1.CH505 w4.3 (purple) in macaques immunized at different times with both Zika prM-E and a series of HIV-1 CH505 gp160 mRNA. ZIKV neutralization was measured by the PRNT assay. **c.** Plasma IgG binding titers (log area-under-the-curve) to HIV-1 CH505 TF gp120 (purple) and Zika prM-E (forest green) for the macaques shown in **b**. The line represents the group mean binding titer.

While these results indicated that durable neutralizing antibodies were elicited by each vaccine regimen, the downward trend of the neutralization titers in these macaques and the A244-immunized macaques raised the question of how long would neutralizing antibodies persist at detectable levels. A subset of the macaques used in this HIV-1 study were previously administered 50 μg of mRNA-LNP encoding ZIKV pre-membrane and envelope (prM-E). These macaques generated protective anti-Zika neutralizing antibody responses ^1^. Using these macaques that were administered both HIV and Zika mRNA-LNP vaccines, we determined serum neutralizing antibodies titers over 52 weeks (**Fig. 7B**). We found that neutralizing antibodies against ZIKV were maintained until the last timepoint of follow-up 52 weeks after immunization. Neutralizing antibodies to HIV-1 persisted for the duration of the 41 weeks of follow-up. During these 41 weeks, HIV-1 titers initially fell approximately 10-fold after the last immunization but plateaued at ~1:100 titer (**Fig. 7B**). In contrast, Zika neutralizing antibody levels were maintained at the same level for the 52 weeks after being detected at week 2 (**Fig. 7B**). Similar patterns were observed for plasma binding IgG titers to CH505 gp120 and Zika prM-E (**Fig. 7C**). Taken together, nucleoside-modified mRNA-LNP immunizations induced durable serum neutralizing antibodies for both HIV-1 and Zika viruses.

### Neutralizing antibody titers are dependent on mRNA-LNP dose

In our previous ZIKV vaccine studies in nonhuman primates, we administered 50, 200 or 600 μg of mRNA-LNP^1^. Given the potent elicitation of neutralizing antibodies by mRNA administered at each of these doses, we sought to determine whether the mRNA dose could be further decreased. We immunized four macaques intramuscularly with either 50, 20, or 5 μg of mRNA-LNP encoding Zika prM-E (**Fig. 8A**) and compared titers of binding IgG and neutralizing antibodies. Fifity and twenty microgram doses of mRNA-LNP elicited similar titers of binding IgG and neutralizing antibodies (**Fig. 8B** and **C**). Titers of ZIKV binding IgG and neutralizing antibodies were substantially decreased when the mRNA-LNP dose was lowered to 5 μg, but were detectable with administration of this single administration of a small amount of mRNA-LNP in macaques. The route of immunization was not critical as administering 50 μg of mRNA-LNP either intramuscularly or intradermally elicited comparable Zika envelope binding IgG and Zika neutralizing antibodies (**Fig. 8B** and **C**). Thus, the mRNA-LNP dose could be lowered from 50 to 20 μg in macaques without detrimental effects to antibody responses. Thus, more immunizations can be performed with each preparation of mRNA-LNP by reducing the dose by 60 percent.

**Figure 8.**
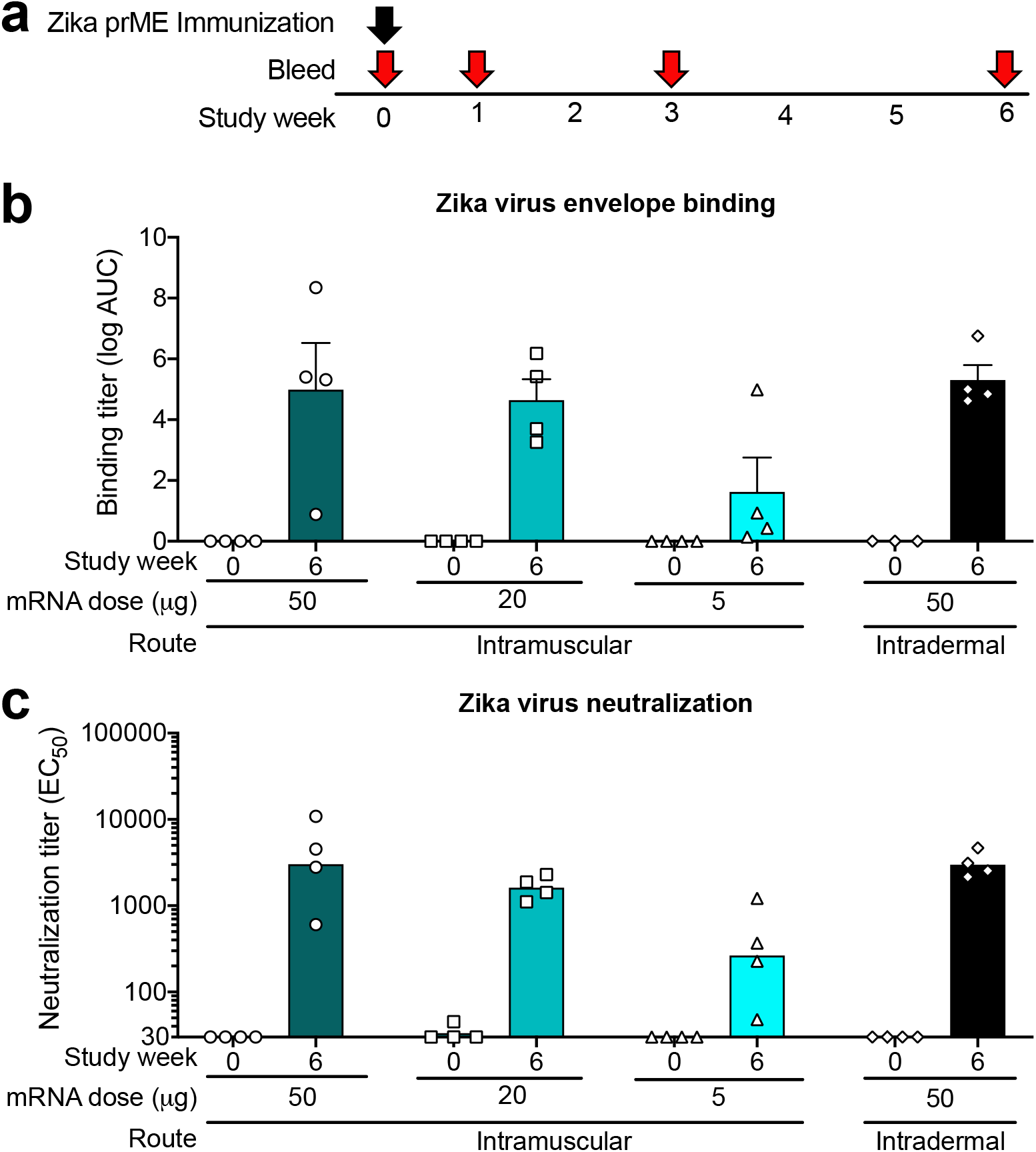
Neutralizing antibody titers are dependent on mRNA dose, but are similar for intramuscular and intradermal immunization routes. **a.** Immunization regimen for groups of 4 rhesus macaques immunized with different amounts of mRNA-LNP via intramuscular or intradermal routes. **b.** Plasma binding IgG titers specific for ZIKV envelope protein were determined by ELISA before vaccination (week 0) or 6 weeks post-vaccination. Binding IgG titers are shown as log area-under-the-curve. Each symbol represents an individual macaque and the bar shows the group mean and standard error. **c.** Serum neutralization titers against ZIKV determined by the reporter virus particle assay are shown as the 50% effective concentration (EC_50_). Symbols are the same as in **b** and bars represent the group geometric mean.

## Discussion

Here, we demonstrate in 28 rhesus macaques that immunization with nucleoside-modified mRNA-LNP (n=16) was equal to or superior to the immunogenicity of adjuvanted Env protein. There are multiple factors for why nucleoside-modified mRNA immunization has potent immunogenicity for viral proteins. First, mRNA is delivered by LNP to dendritic cells and likely other immune cells that are able to activate naïve T cells to respond to the vaccine immunogen ^59–61^. Second, nucleoside-modified mRNA-LNP immunization generates robust antigen-specific germinal center T and B cell responses ^2^. Within the germinal center, follicular T helper cells provide help to B cells undergoing affinity maturation ^34^. The affinity maturation process is critical for the development of high affinity antibodies after vaccination, and in particular for HIV-1 is required for bnAb development ^62^. Animal models vaccinated with nucleoside-modified mRNA-LNP encoding HIV-1 Env or influenza virus hemagglutinin develop high levels of pathogen-specific T follicular helper cells ^33^. Given that HIV-1 neutralizing antibody breadth is correlated with the frequency of circulating T follicular helper cells ^35,63^, nucleoside-modified mRNA-LNP immunization warrants further study as an HIV-1 vaccine platform. Nucleoside-modified mRNAs have many potential advantages over proteins including speed of production and cost-savings. Moreover, mRNA-LNP increases the feasibility of making under current good manufacturing practices the multicomponent vaccines that are needed to induce both protective HIV-1 bnAbs and nnAbs.

In RV144, aluminum hydroxide (Rehydragel) was used and poor durability of protective immune responses was observed ^24,64^. ALFQ is a potent adjuvant that contains anionic liposomes, QS21, and TLR-4 agonist, MPL ^39^, both potent adjuvants. Thus, comparison of mRNA-LNP with Rehydragel gives a direct indication of how it might fare versus the RV144 regimen, and comparison with ALFQ provides an indication of how it might fare in comparison with a stronger adjuvant. The adjuvant GLA (synthetic MPL) in SE (stable emulsion) ^58^ has been used in HVTN 115 with multiple gp120s and was immunogenic, although in a head-to-head comparison of Rehydragel and GLA-SE in primates, they were similar in promoting immunogenicity except for V2 responses where Rehydragel was superior. While ALFQ and GLA stable emulsion contains lipids, they differ from the lipid nanoparticles. The lipid nanoparticles include ionizable cationic lipids that give the lipid nanoparticle a neutral surface charge, whereas most commercial liposomes lack ionizable cationic lipids. The neutral charge helps reduce non-specific protein binding in vivo. Unlike GLA or ALFQ, the LNP does not activate TLR-4, but instead activates follicular helper T cells^33^. In general, with the doses and adjuvants used in this study of proteins versus mRNA-LNP, nucleoside-modified mRNA-LNP immunization was superior or equal to protein in Rehydragel or GLA-SE and equal to protein in ALFQ. Future studies using different vaccination intervals, adjuvants, and protein doses could corroborate this conclusion.

There are currently two ongoing efficacy trials that test the protective capacity of non-neutralizing HIV-1 antibodies with viral vector prime, Env protein boosts to determine if the 42 month efficacy of ALVAC/gp120 can be replicated. In addition, the just-completed HVTN 702 trial (NCT02968849) that uses an ALVAC with a clade C Env insert as prime and ALVAC-C + a bivalent C gp120 Env boost failed to confer any protection from HIV-1 infection in South Africa in an area of high HIV-1 infection rates. The ongoing HVTN 705 trial (NCT03060629) tests the efficacy of an adenovirus (Ad) 26 vector containing mosaic HIV-1 genes as a prime and the same Ad26 + a clade C gp140 Env as a boost, and the HVTN 706 trial (NCT03964415) tests the efficacy of Ad26 mosaic HIV as a prime and Ad26 + a bivalent clade C gp140 Env + a mosaic gp140 Env as boost ^65^. The correlate of decreased transmission risk in macaque studies of the A26, Ad26 + gp140 boost SHIV challenge studies did not include neutralization breadth of antibodies, but instead included clade C Env ELISA binding antibodies and interferon gamma Env ELISPOT values ^65^. Thus, these latter two vaccines will test whether non-neutralizing antibody effector functions can mediate significant protection in humans.

The ease and cost-effectiveness of mRNA-LNP production compared to the production of multiple recombinant proteins by good manufacturing practice (GMP) techniques makes mRNA-LNP particularly attractive. Furthermore, the scalability of mRNA-LNP for thousands to millions of doses again make the cost and ease of production of mRNA-LNP attractive. These manufacturing considerations are key for improving on RV144, and show great promise for COVID-19 vaccines currently under development ^17^. However, the cold temperature required for distribution and storage of mRNA vaccines remains a challenge for this vaccine platform ^66^.

While mRNA vaccination is promising, there are still improvements that can be made for HIV mRNA-LNP vaccination. Specifically, for bnAb induction by mRNAs, the near-native Env trimers will need to be stabilized such that they will be able to be produced by the transfected cell in a well-folded state. This challenge is not unique to mRNA as DNA vaccination has also faced similar roadblocks^67^. This aspect of genetic vaccination differs from recombinant proteins that can be purified prior to immunization and recognition by the immune system. However, there is a plethora of available mutations for stabilizing HIV-1 envelope, and each envelope has different intrinsic stability that can be further augmented. We investigated only two potential sets and found H66A and A587T reduced non-neutralizing antibody binding to envelope in vitro. The reduction in non-neutralizing antibody binding suggests a higher percentage of native-like envelope expressed by the mRNA. Future vaccine studies will need to determine whether reduced non-neutralizing antibody binding in vitro translates to improved neutralizing antibody elicitation in vivo. Despite the best envelope stabilization designs, it is possible that not all of the envelope will be well-folded, native-like trimers. In vivo studies by us and others are aiming to determine what the percentage of well-folded envelope needs to be in order to elicit optimal neutralizing antibody responses. Both of these important questions are areas of intense investigation and are critical for mRNA becoming a widely accepted HIV-1 vaccine platform.

In summary, comparison of nucleoside-modified mRNA-LNP with recombinant Env proteins in three adjuvants demonstrated the utility of mRNA-LNP as a mode of inducing nnAbs that are predicted to be protective against retrovirus challenge. We suggest that moving to nucleoside-modified mRNA-LNP for the next generation of clinical trials for multivalent Env immunization will be advantageous and speed the development of a globally available protective HIV-1 vaccine.

## Acknowledgements

We acknowledge technical assistance from Haiyan Chen, Esther Lee, Kedamawit Tilahun, Andrew Foulger, Aja Sanzone. This work was supported NIH Division of AIDS UM1 grant AI100645 for the Center for HIV/AIDS Vaccine Immunology-Immunogen Discovery (CHAVI-ID; B.F.H.), NIH NIAID Duke Center for AIDS Research grant P30 AI064518 (S.X.S., G.D.T), and the intramural program of NIAID. The funders had no role in data collection and interpretation, or the decision to submit the work for publication.

## Author contributions

Experimental Design: KOS, NP, DW, BFH; Investigation and assays: NP, RP, LS, RS, MB, AE, GH, XS, DG, DNG, CB, YT, ZM; Supervision: KOS, SS, ML, TCP, XS, GF, GT, DCM, DW, BFH; Data analysis: KOS, NP, RP, MB, AE, ML, RWR, YW, DNG, TCP, XS, GF, GT, DCM, DW, BFH; Funding: XS, TCP, GT, BFH; Wrote manuscript: KOS, NP, DW, BFH.

## Declaration of Interests

The authors declare no financial conflicts of interest. B.F.H. and K.O.S. have patent applications submitted on CH505 Envs used in this study. T.C.P. is an inventor on multiple patent applications related to ZIKV vaccine antigen constructs and vaccines. In accordance with the University of Pennsylvania policies and procedures and our ethical obligations as researchers, we report that D.W. is named on patents that describe the use of nucleoside-modified mRNA as a platform to deliver therapeutic proteins. D.W. and N.P. are also named on a patent describing the use of nucleoside-modified mRNA in lipid nanoparticles as a vaccine platform. We have disclosed those interests fully to the University of Pennsylvania, and we have in place an approved plan for managing any potential conflicts arising from licensing of our patents.

## Methods

### Animals and Immunizations

Rhesus macaques were housed and treated in AAALAC-accredited institutions. The study protocol and all veterinarian procedures were approved by the Duke University IACUC and were performed based on standard operating procedures. Macaques were anesthetized with ketamine for immunizations and blood collection. For protein immunizations, rhesus macaques were immunized with 300 μg of A244 gp120 monomers in the quadriceps. In previous macaque studies 300 μg and 100 μg of A244 gp120 monomers gave equivalent antibody responses. Thus, 100 μg of HIV-1 CH505 stabilized SOSIP gp140 recombinant protein were administered in the quadriceps of rhesus macaques instead of 300 μg. Total immunization volume was 1 mL of protein and adjuvant diluted in phosphate-buffered saline (PBS). For RNA immunizations, rhesus macaques were shaved on their backs and injected intradermally with nucleoside-modified mRNA-LNP diluted in PBS. For ZIKV immunization, three macaques received 600 μg and two macaques received 200 μg of mRNA expressing Zika preM-E ^1^. In our previously published mRNA-LNP titration, there was no difference in antibody titers with 600, 200, and 50 μg of mRNA-LNP expressing Zika preM-E^1^; thus, for HIV-1 immunizations we administered 50 μg of mRNA-LNP expressing HIV-1 Env. Blood was collected by femoral venipuncture under ketamine anesthesia. Serum and EDTA-plasma were isolated from whole blood and stored at −80°C ^1^.

### Production of Recombinant Env protein immunogens

Recombinant HIV-1 Env gp120 was expressed in Freestyle 293 cells (ThermoFisher) by transient transfection. To transfect Freestyle 293 cells, 1 milligram of plasmid DNA encoding HIV-1 Env per 1 liter of cells was diluted in DMEM and mixed with polyethyleneimine (PEI, Polysciences, Inc.). The DNA was allowed to complex with PEI for 25 min and then cells were added. The Freestyle 293 cells were incubated with the PEI:DNA mixtures for 4 h. Freestyle 293 (ThermoFisher) cells were subsequently centrifuged and resuspended in Freestyle293 media (ThermoFisher) at a final density of 1.25 million cells/mL. Four days after transfection, the cell culture media was cleared of cells by centrifugation and filtered with 0.8 μm filter. A Vivaflow 50 with a 10 kD MWCO (ThermoFisher) was used to concentrate the cell-free culture media. The concentrated cell culture supernatant was incubated with Galanthus Nivalis lectin beads (Vistar Laboratories) rotating overnight at 4 °C. The beads were pelleted by centrifugation the next day and resuspended in MES wash buffer (20mM MES, 10mM CaCl_2_, 130mM NaCl pH7.0). The lectin beads were washed twice with MES wash buffer, and the protein was eluted with MES wash buffer supplemented with 500 mM methyl-alpha-D-mannopyranoside (Sigma). The protein was buffer exchanged into phosphate buffered saline by successive rounds of centrifugation and stored at −80°C.

HIV-1 Env soluble trimers were designed as gp41-chimeric SOSIP gp140s ^68–70^. The gp41-chimeric SOSIP gp140s were further stabilized by the addition of E64K and A316W mutations ^68^. CH505 SOSIP gp140 Envs were expressed as previously described ^71^. Freestyle293 cells (Invitrogen) were diluted to 1.25×10^6^ cells/mL with fresh Freestyle293 media up to 1L total volume. The cells were co-transfected with 650 μg of SOSIP expressing plasmid DNA and 150 μg of furin expressing plasmid DNA complexed with 293Fectin (Invitrogen). The cell culture supernatant clarified on day 6 and subjected to positive PGT145 antibody affinity chromatography. Eluted protein was concentrated down to 2 mL for size exclusion chromatography with a Superose6 16/600 column (GE Healthcare) in 10 mM Tris pH8, 500 mM NaCl. Fractions containing trimeric HIV-1 Env protein were pooled together, sterile-filtered, snap frozen, and stored at −80°C.

### mRNA production

mRNAs were produced as previously described ^72^ using T7 RNA polymerase (Megascript, Ambion) on linearized plasmids encoding codon-optimized ^73^ ZIKV prM-E (pUC-TEV-ZIKV prM-E-A101) from the strain H/PF/2013 (Asian lineage, French Polynesia, 2013, GenBank: KJ776791); gp160 HIV-1 Env: coCH505.M5 Env (pUC-TEV-coCH505.M5 Env-A101), coCH505.M11 Env (pUC-TEV-coCH505.M11 Env-A101), coCH505w0.30.20 Env (pUC-TEV-coCH505w0.30.20 Env-A101), coCH505w0.136.B18 Env (pUC-TEV-coCH505w0.136.B18 Env-A101), coCH505w0.20.14 Env (pUC-TEV-coCH505w0.20.14 Env-A101), coCH505w0.30.12 Env (pUC-TEV-coCH505w0.30.12 Env-A101); chimeric stabilized HIV-1 Envs: CH505M5chim.6R.SOSIP.664v4.1 Env (pUC-TEV-CH505M5chim.6R.SOSIP.664v4.1 Env-A101), CH505M11chim.6R.SOSIP.664v4.1 Env (pUC-TEV-CH505M11chim.6R.SOSIP.664v4.1 Env-A101), CH505w20.14chim.6R.SOSIP.664v4.1 Env (pUC-TEV-CH505w20.14chim.6R.SOSIP.664v4.1 Env-A101), CH505w30.12chim.6R.SOSIP.664v4.1 Env (pUC-TEV-CH505w30.12chim.6R.SOSIP.664v4.1 Env-A101), CH505w30.20chim.6R.SOSIP.664v4.1 Env (pUC-TEV-CH505w30.20chim.6R.SOSIP.664v4.1 Env-A101), CH505w136.B18chim.6R.SOSIP.664v4.1 Env (pUC-TEV-CH505w136.B18chim.6R.SOSIP.664v4.1 Env-A101); gp120 HIV-1 Envs: A244delta11 gp120 Env (pUC-TEV-A244delta11 gp120 Env-A101), A244delta11 K368R gp120 Env (pUC-TEV-A244delta11 K368R gp120 Env-A101). mRNAs were transcribed to contain 101 nucleotide-long poly(A) tails based on the DNA-encoded poly(A) tail sequence. One-methylpseudouridine (m1Ψ)-5’-triphosphate (TriLink) instead of UTP was used to generate modified nucleoside-containing mRNA. RNAs were capped using the m7G capping kit with 2’-*O*-methyltransferase (ScriptCap, CellScript) to obtain a type 1 cap. mRNA was purified by Fast Protein Liquid Chromatography (FPLC) (Akta Purifier, GE Healthcare), as described ^74^. All mRNAs were analyzed by denaturing or native agarose gel electrophoresis and were stored frozen at −20°C.

### LNP formulation of mRNA

FPLC-purified m1Ψ-containing HIV-1 Env and ZIKV prM-E mRNAs were encapsulated in LNP using a self-assembly process in which an aqueous solution of mRNA at pH=4.0 is rapidly mixed with a solution of lipids dissolved in ethanol ^75^. LNP used in this study were similar in composition to those described previously ^75,76^, which contain an ionizable cationic lipid (proprietary to Acuitas)/phosphatidylcholine/cholesterol/PEG-lipid. The proprietary lipid and LNP composition are described in US patent US10,221,127. They had a diameter of ~80 nm as measured by dynamic light scattering using a Zetasizer Nano ZS (Malvern Instruments Ltd, Malvern, UK) instrument.

### *In vitro* mRNA transfection and flow cytometry

*In vitro* mRNA transfection and antibody binding were performed as earlier described^77^. Freestyle 293F cells were maintained at 37°C with 8% CO_2_ and shaking at 130 rpm. Cells were diluted to 1×10^6^ cells/ml prior to transfection. Bare mRNA encoding CH505 M5 gp160 Envs were transfected at 0.5 μg/million cells using TransIT-mRNA Transfection Kit (Mirus). Transfected cells were harvested 48 hr post transfection. Cells were first incubated with HIV antibodies at 10 μg/ml for 30 min at 4°C in dark. Antibody binding was then detected by Goat F(ab’)_2_ Anti-Human IgG - (Fab’)_2_ (PE) (used at 2.5 μg/ml) by incubating cells with the secondary antibody for 30 min at 4°C in dark. Finally, data were collected on a LSRII with a high-throughput system.

### HIV-1 Env direct ELISAs

Direct ELISAs were performed as stated previously ^78,79^. Specifically, NuncSorp 384-well plates were coated with 30 ng (2 μg/mL) of antigen in 0.1 M sodium bicarbonate overnight at 4°C. Plates were washed with SuperWash and blocked with SuperBlock. Plates were washed. Plasma was serially diluted in SuperBlock and added to the ELISA plate. The plate was washed twice and a 1:8000 dilution of anti-rhesus IgG conjugated to horseradish peroxidase (HRP; Rockland Immunochemicals) was incubated in the plate for 1 h. The plate was washed and tetramethylbenzidine (TMB; KPL) was incubated in the plate for 15 min to detect secondary antibody binding. The reaction was stopped with 1% HCl and absorbance at 450 nm was read with a SpectraMax plate reader (Molecular Devices). SoftMax Pro v5.3 was used to calculate log area-under-the-curve (AUC; Molecular Devices).

### Plasma competition ELISAs

Plasma competition assays were performed as described previously ^78,79^. In brief, NuncSorp plates were coated with HIV-1 Env, washed and blocked as stated above for direct ELISAs. After blocking was complete, nonhuman primate plasma was diluted in SuperBlock at a 1:50 dilution and incubated in triplicate wells for 90 min. Non-biotinylated monoclonal antibodies were incubated with the Env in triplicate as positive controls for blocking. To determine relative binding no plasma or no antibody was added to a group of wells scattered throughout the plate. After 90 min the non-biotinylated antibody or plasma was washed away and biotinylated monoclonal antibodies was incubated in the wells for 1 h at subsaturating concentrations. Specifically, biotinylated monoclonal antibody concentrations used for binding to A244 Δ11 gp120 were: 0.04 μg/mL of CH58, 0.1 μg/mL of A32, 1.9 μg/mL of 2G12, 0.2 μg/mL of PGT125, 1.2 μg/mL of CH01, 1μg/mL of PG9. Biotinylated CH01 and PG9 concentrations used for binding to 9021 gp140 and B.6240 Δ11 gp120 were 1.5 and 0.125 μg/mL respectively. For CD4 blocking assays, soluble CD4 was added followed by biotinylated anti-CD4 monoclonal antibody. Each well was washed and binding of biotinylated monoclonal antibodies was determined with a 1:30000 dilution of HRP-conjugated streptavidin (ThermoFisher). HRP was detected with TMB and stopped with 1% HCl. The absorbance at 450 nm of each well was read with a Spectramax plate reader (Molecular Devices). Binding of the biotinylated monoclonal antibody to HIV-1 Env in the absence of plasma was compared to in the presence of plasma to calculate percent inhibition of binding. Based on historical negative controls, assays were considered valid if the positive control antibodies blocked greater than 20% of the biotinylated antibody binding.

### HIV-1 Env Peptide Array

Peptide arrays were performed as previously described ^80^. The HIV-1 peptide libraries contain overlapping HIV-1 peptides covering full-length gp120 of 5 consensus viruses from group M and clades A, B, C, and D ^80^. Array slides were provided by JPT Peptide Technologies GmbH (Germany) by printing a library of peptides onto epoxy glass slides (PolyAn GmbH, Germany). The library contains overlapping peptides (15-mers overlapping by 12) covering 5 full-length gp160 consensus sequences (clade A, B, C, D, and group M). V3 peptide binding breadth was analyzed for a library of V3 peptides (15-mers overlapping by 12) for 7 consensus sequences (clade A, B, C, D, CRF1, and CRF2, and group M) and 6 vaccine strains (MN, A244, TH023, TV-1, ZM651, 1086C). Three identical subarrays were blocked for 1 h, followed by a 2-h incubation with monoclonal antibody, and a subsequent 45-min incubation with anti-monkey IgG conjugated with AF647 (Jackson ImmunoResearch, PA). Slides were washed before the addition of monoclonal antibody or anti-monkey IgG. Unbound anti-monkey IgG conjugated with AF647 was washed away, and array slides were scanned at a wavelength of 635 nm using an InnoScan 710 scanner (InnopSys, Denmark) and images were analyzed using Magpix V8.1.1.

### Antibody-Dependent Cellular Cytotoxicity (ADCC)

ADCC activity was measured by ADCC GranToxiLux (GTL) ^81^ and tested against subtype AE HIV-1 recombinant A244 Δ11 gp120-coated cells. NHP plasma were incubated with human PBMC as source of effector cells ^82^ and gp120-coated target CEM.NKR._CCR5_ cells _83_ and ADCC was quantified as net percent granzyme B activity, which is the percent of target cells positive for GranToxiLux (GTL) detected by flow cytometry. For each subject at each timepoint, percent granzyme B activity was measured at six dilution levels: 50, 250, 1250, 6250, 31,250 and 156,250 for each antigen. Peak activity less than 0% was set to 0%. A positive response was defined as peak activity greater than or equal to 8%. The RSV-specific monoclonal antibody Palivizumab and a cocktail of HIV-1 monoclonal Abs (A32, 2G12, CH44, and 7B2) were used as negative and positive control, respectively. The potency of ADCC responses was evaluated by calculating the Area Under the Curve using the non-parametric trapezoidal method.

### Infected Cells Ab Binding Assay (ICABA)

The measurement of plasma Ab binding to HIV-1 Env expressed on the surface of infected cells was conducted using flow-cytometry-based indirect surface staining according to methods similar to those previously described ^41,84^. Briefly, mock infected and the subtype AE HIV-1 CM235-infected (HIV-CM235-2.LucR.T2A/293T/17, Accession Number AF259954.1) CEM.NKR_CCR5_ cells _83_ were incubated with 1:100 dilutions of plasma samples collected from the animals before immunizations and 2 weeks after the 2^nd^ immunization (week 8) for 2h at 37°C, then stained with a vital dye (Live/Dead Aqua) to exclude dead cells from analysis. The cells were subsequently washed, and permeabilized using BD Cytofix/Cytoperm solution. After an additional wash, cells were stained with FITC-conjugated goat-anti-Rhesus polyclonal antisera (Southern Biotech) to detect binding of the plasma Ab (pAb), and RD1-conjugated anti-HIV-1 p24 (KC57, Beckman Coulter) to identify infected cells. Backgroud signal was defined as signal obtained when staining cells with the secondary antibody alone, and signal observed when staining the mock-infected cells. Cells scored as positive for NHP plasma binding were defined as viable, p24 positive, and FITC positive. Final results are reported as the percent of FITC positive cells (p24+pAb+) and FITC MFI among the viable p24 positive events after subtracting the background. Baseline correction for the FITC MFI was performed by subtracting pre-immunization MFI wherever applicable.

### Antibody-dependent phagocytosis

Phagocytosis was measured as stated previously ^85,86^. Briefly, recombinant A244 Δ11 gp120 was incubated with 1 μM fluorescent beads overnight at 4°C while rotating. The beads were subsequently washed twice with 0.1% BSA/PBS to remove unbound gp120. gp120-coated beads were incubated with prevaccination and postvaccination plasma for 2 h at 37°C. As a positive control HIVIG was incubated with the beads. As a negative control monoclonal antibody CH65 was incubated with the beads, since this antibody reacts with influenza hemagglutinin and not HIV-1 Env. THP-1 or monocytes were incubated with anti-CD4 antibody SK3 (Biolegend) and then added to the bead/plasma mixture. The mixture was spinoculated for 1h at 4C followed by another 1h incubation at 37°C ^87^. The cells were washed and fixed and fluorescence due to bead internalization was measured by flow cytometry. Positive phagocytosis was determined by two criteria. First, values at week 84 were considered positive if they were 3 standard deviations above values obtained in the prevaccination sample from the same macaque. Second, values obtained at week 84 had to be >3-fold higher than the prevaccination sample from the same macaque.

### HIV-1 neutralization

Neutralization assays were performed in TZM-bl cells using 293T-produced Env-pseudotyped viruses in a 96-well format as described ^88^. Neutralization titers are the serum dilution at which relative luminescence units (RLUs) plotted on a non-linear regression curve were reduced by 50% compared to virus control wells after subtraction of background RLUs in uninfected cell control wells.

### ELISA binding of plasma IgG to Zika E protein

ELISA binding was performed as described previously ^89^. ZIKV envelope or HIV-1 proteins were coated overnight at 4°C on the wells of 384-well plates (Corning Life Sciences). Each protein was coated at 2 μg/mL in 15 μl in sodium bicarbonate. Each well of the plate was blocked with assay diluent (PBS containing 4% [wt/vol] whey protein–15% normal goat serum–0.5% Tween 20–0.05% sodium azide) for 2 h at room temperature. Blocking solution was aspirated from each well and 10 μL of serially-diluted macaque sera were added to the plate and incubated for 90 min. The plate was washed with PBS-Tween-20. Ten microliters of horse radish peroxidase anti-macaque IgG Fc was added to each well of the plate and incubated for 1 h. Subsequently, the plate was washed with PBS-Tween-20 and 30 μL of SureBlue substrate was used to visualize the presence of horse radish peroxidase. After 5-15 min the reaction was stopped with 15 μL of 0.1M HCl. Absorbance at 405nm was read for each well using a VersaMax microplate reader (Molecular Devices). Binding titers observed for the prevaccination timepoint were subtracted from the post vaccination timepoint for macaques that were part of mRNA-LNP dose study.

### ZIKV MR-766 plaque reduction neutralization tests (PRNT)

Serum neutralization was assessed as detailed previously ^1^. Fifty plaque forming units of ZIKV strain MR-766 (African lineage, Uganda, 1947, GenBank: AY632535) (UTMB Arbovirus Reference Collection) were incubated with increasing dilutions of heat-inactivated sera in serum-free DMEM (Corning) medium for 1 h at 37°C. The virus/serum mixture was added to a confluent monolayer of Vero cells for 1.5 h at 37°C with intermittent rocking. Three ml of overlay, containing a final concentration of 0.5% methylcellulose (4,000 centipoise) (Sigma), 1X DMEM (Gibco), 16 mM HEPES, 0.56% sodium bicarbonate, 1.6X GlutaMAX (Gibco), 1X penicillin/streptomycin (Corning), and 4 μg/ml amphotericin B (Gibco), was added to each well. Plates were incubated for 5 days at 37°C in 5% CO_2_. After incubation, the overlay was aspirated, cells were fixed and stained with 0.5% crystal violet (Sigma) in 25% methanol, 75% deionized water, rinsed with deionized water, and plaques inspected. Neutralization titers (EC_50_) were calculated as the reciprocal dilution of sera required for 50% neutralization of infection. EC_50_ titers below the limit of detection are reported as half of the limit of detection.

### Zika reporter virus particle (RVP) neutralization assay

RVP neutralization assays were performed as described elsewhere^1^. RVPs were diluted to ensure antibody excess at informative portions of the antibody dose response curve and incubated for 1 h at 37°C with serial dilutions of heat-inactivated macaque sera to allow for steady-state binding. Serum-RVP mixtures were subsequently mixed with Raji-DCSIGNR cells at 37°C. Each serum sample was tested in to technical replicates. Every assay was repeated in two biologically-independent assays. GFP-positive infected cells were detected by flow cytometry 24–48 hr later. The EC50 was estimated using a non-linear regression with a variable slope (GraphPad Prism). The initial dilution of sera (based on the final volume of RVPs, cells, and sera) was set as the limit of quantification of the assay. EC50 titers below the limit of quantification were reported as a titer half the limit of quantification.

## Statistical analyses

Descriptive statistics were calculated and reported for each assay. Group means, standard error, and geometric means were calculated with GraphPad Prism 8. Log area-under-the-curve (AUC) was calculated by Softmax Pro. EC50 titers for RVP neutralization were analyzed by non-linear regression to estimate the reciprocal dilution of sera required for half-maximal neutralization of infection (EC50 titer). The geometric mean of duplicate RVP neutralization EC50 titers were calculated using GraphPad Prism 8.

## Data availability

The authors declare that the data supporting the findings of this study are available within the main and supplemental figures. All data is available from the corresponding author upon reasonable request.

## Notes

### Competing Interest Statement

The authors have declared no competing interest.

## References

1 Pardi, N. et al. Zika virus protection by a single low-dose nucleoside-modified mRNA vaccination. Nature 543, 248–251, doi:10.1038/nature21428 (2017).

2 Alameh, M. G., Weissman, D. & Pardi, N. Messenger RNA-Based Vaccines Against Infectious Diseases. Current topics in microbiology and immunology, doi:10.1007/82_2020_202 (2020).

3 Awasthi, S. et al. Nucleoside-modified mRNA encoding HSV-2 glycoproteins C, D, and E prevents clinical and subclinical genital herpes. Science immunology 4, doi:10.1126/sciimmunol.aaw7083 (2019).

4 Freyn, A. W. et al. A Multi-Targeting, Nucleoside-Modified mRNA Influenza Virus Vaccine Provides Broad Protection in Mice. Molecular therapy: the journal of the American Society of Gene Therapy, doi:10.1016/j.ymthe.2020.04.018 (2020).

5 Lo, M. K. et al. Evaluation of a Single-Dose Nucleoside-Modified Messenger RNA Vaccine Encoding Hendra Virus-Soluble Glycoprotein Against Lethal Nipah virus Challenge in Syrian Hamsters. The Journal of infectious diseases 221, S493–s498, doi:10.1093/infdis/jiz553 (2020).

6 Nelson, C. S. et al. Human Cytomegalovirus Glycoprotein B Nucleoside-Modified mRNA Vaccine Elicits Antibody Responses with Greater Durability and Breadth than MF59-Adjuvanted gB Protein Immunization. Journal of virology 94, doi:10.1128/jvi.00186-20 (2020).

7 Pardi, N., Hogan, M. J. & Weissman, D. Recent advances in mRNA vaccine technology. Current opinion in immunology 65, 14–20, doi:10.1016/j.coi.2020.01.008 (2020).

8 Pardi, N., Hogan, M. J., Porter, F. W. & Weissman, D. mRNA vaccines - a new era in vaccinology. Nature reviews. Drug discovery 17, 261–279, doi:10.1038/nrd.2017.243 (2018).

9 Kariko, K., Buckstein, M., Ni, H. & Weissman, D. Suppression of RNA recognition by Toll-like receptors: the impact of nucleoside modification and the evolutionary origin of RNA. Immunity 23, 165–175, doi:10.1016/j.immuni.2005.06.008 (2005).

10 Kariko, K. et al. Incorporation of pseudouridine into mRNA yields superior nonimmunogenic vector with increased translational capacity and biological stability. Molecular therapy: the journal of the American Society of Gene Therapy 16, 1833–1840, doi:10.1038/mt.2008.200 (2008).

11 Gustafsson, C., Govindarajan, S. & Minshull, J. Codon bias and heterologous protein expression. Trends in biotechnology 22, 346–353, doi:10.1016/j.tibtech.2004.04.006 (2004).

12 Kariko, K., Muramatsu, H., Ludwig, J. & Weissman, D. Generating the optimal mRNA for therapy: HPLC purification eliminates immune activation and improves translation of nucleoside-modified, protein-encoding mRNA. Nucleic Acids Res 39, e142, doi:10.1093/nar/gkr695 (2011).

13 Pardi, N. et al. Expression kinetics of nucleoside-modified mRNA delivered in lipid nanoparticles to mice by various routes. J Control Release 217, 345–351, doi:10.1016/j.jconrel.2015.08.007 (2015).

14 Akinc, A. et al. The Onpattro story and the clinical translation of nanomedicines containing nucleic acid-based drugs. Nature nanotechnology 14, 1084–1087, doi:10.1038/s41565-019-0591-y (2019).

15 Sempowski, G. D., Saunders, K. O., Acharya, P., Wiehe, K. J. & Haynes, B. F. Pandemic Preparedness: Developing Vaccines and Therapeutic Antibodies For COVID-19. Cell, doi:10.1016/j.cell.2020.05.041 (2020).

16 Jackson, L. A. et al. An mRNA Vaccine against SARS-CoV-2 - Preliminary Report. N Engl J Med, doi:10.1056/NEJMoa2022483 (2020).

17 Mulligan, M. J. et al. Phase 1/2 study of COVID-19 RNA vaccine BNT162b1 in adults. Nature, doi:10.1038/s41586-020-2639-4 (2020).

18 Keech, C. et al. Phase 1-2 Trial of a SARS-CoV-2 Recombinant Spike Protein Nanoparticle Vaccine. N Engl J Med, doi:10.1056/NEJMoa2026920 (2020).

19 Haynes, B. F., Burton, D. R. & Mascola, J. R. Multiple roles for HIV broadly neutralizing antibodies. Science translational medicine 11, doi:10.1126/scitranslmed.aaz2686 (2019).

20 Haynes, B. F., Kelsoe, G., Harrison, S. C. & Kepler, T. B. B-cell-lineage immunogen design in vaccine development with HIV-1 as a case study. Nature biotechnology 30, 423–433, doi:10.1038/nbt.2197 (2012).

21 Zhao, L. P. et al. Landscapes of binding antibody and T-cell responses to pox-protein HIV vaccines in Thais and South Africans. PloS one 15, e0226803, doi:10.1371/journal.pone.0226803 (2020).

22 Cohen, J. Another HIV vaccine strategy fails in large-scale study. Science (New York, N.Y.), doi:doi:10.1126/science.abb1480 (2020).

23 Rerks-Ngarm, S. et al. Vaccination with ALVAC and AIDSVAX to prevent HIV-1 infection in Thailand. The New England journal of medicine 361, 2209–2220, doi:10.1056/NEJMoa0908492 (2009).

24 Haynes, B. F. et al. Immune-correlates analysis of an HIV-1 vaccine efficacy trial. The New England journal of medicine 366, 1275–1286, doi:10.1056/NEJMoa1113425 (2012).

25 Seabright, G. E., Doores, K. J., Burton, D. R. & Crispin, M. Protein and Glycan Mimicry in HIV Vaccine Design. Journal of molecular biology 431, 2223–2247, doi:10.1016/j.jmb.2019.04.016 (2019).

26 Kwong, P. D. et al. HIV-1 evades antibody-mediated neutralization through conformational masking of receptor-binding sites. Nature 420, 678–682, doi:10.1038/nature01188 (2002).

27 Wei, X. et al. Antibody neutralization and escape by HIV-1. Nature 422, 307–312, doi:10.1038/nature01470 (2003).

28 Haynes, B. F. et al. Cardiolipin polyspecific autoreactivity in two broadly neutralizing HIV-1 antibodies. Science (New York, N.Y.) 308, 1906–1908, doi:10.1126/science.1111781 (2005).

29 Haynes, B. F. & Verkoczy, L. AIDS/HIV. Host controls of HIV neutralizing antibodies. Science (New York, N.Y.) 344, 588–589, doi:10.1126/science.1254990 (2014).

30 Bonsignori, M. et al. Staged induction of HIV-1 glycan-dependent broadly neutralizing antibodies. Science translational medicine 9, doi:10.1126/scitranslmed.aai7514 (2017).

31 Saunders, K. O. et al. Targeted selection of HIV-specific antibody mutations by engineering B cell maturation. Science (New York, N.Y.) 366, doi:10.1126/science.aay7199 (2019).

32 Wiehe, K. et al. Functional Relevance of Improbable Antibody Mutations for HIV Broadly Neutralizing Antibody Development. Cell host & microbe 23, 759–765.e756, doi:10.1016/j.chom.2018.04.018 (2018).

33 Pardi, N. et al. Nucleoside-modified mRNA vaccines induce potent T follicular helper and germinal center B cell responses. The Journal of experimental medicine 215, 1571–1588, doi:10.1084/jem.20171450 (2018).

34 Crotty, S. T follicular helper cell differentiation, function, and roles in disease. Immunity 41, 529–542, doi:10.1016/j.immuni.2014.10.004 (2014).

35 Moody, M. A. et al. Immune perturbations in HIV-1-infected individuals who make broadly neutralizing antibodies. Science immunology 1, aag0851, doi:10.1126/sciimmunol.aag0851 (2016).

36 Locci, M. et al. Human circulating PD-1+CXCR3-CXCR5+ memory Tfh cells are highly functional and correlate with broadly neutralizing HIV antibody responses. Immunity 39, 758–769, doi:10.1016/j.immuni.2013.08.031 (2013).

37 Yates, N. L. et al. HIV-1 Envelope Glycoproteins from Diverse Clades Differentiate Antibody Responses and Durability among Vaccinees. J Virol 92, doi:10.1128/JVI.01843-17 (2018).

38 Alam, S. M. et al. Antigenicity and immunogenicity of RV144 vaccine AIDSVAX clade E envelope immunogen is enhanced by a gp120 N-terminal deletion. Journal of virology 87, 1554–1568, doi:10.1128/jvi.00718-12 (2013).

39 Alving, C. R., Peachman, K. K., Matyas, G. R., Rao, M. & Beck, Z. Army Liposome Formulation (ALF) family of vaccine adjuvants. Expert review of vaccines 19, 279–292, doi:10.1080/14760584.2020.1745636 (2020).

40 Barouch, D. H. et al. Vaccine protection against acquisition of neutralization-resistant SIV challenges in rhesus monkeys. Nature 482, 89–93, doi:10.1038/nature10766 (2012).

41 Bradley, T. et al. Pentavalent HIV-1 vaccine protects against simian-human immunodeficiency virus challenge. Nature communications 8, 15711, doi:10.1038/ncomms15711 (2017).

42 Rolland, M. et al. Increased HIV-1 vaccine efficacy against viruses with genetic signatures in Env V2. Nature 490, 417–420, doi:10.1038/nature11519 (2012).

43 Zolla-Pazner, S. et al. Vaccine-induced IgG antibodies to V1V2 regions of multiple HIV-1 subtypes correlate with decreased risk of HIV-1 infection. PloS one 9, e87572, doi:10.1371/journal.pone.0087572 (2014).

44 Liao, H. X. et al. Vaccine induction of antibodies against a structurally heterogeneous site of immune pressure within HIV-1 envelope protein variable regions 1 and 2. Immunity 38, 176–186, doi:10.1016/j.immuni.2012.11.011 (2013).

45 Pollara, J. et al. Epitope specificity of human immunodeficiency virus-1 antibody dependent cellular cytotoxicity [ADCC] responses. Current HIV research 11, 378–387, doi:10.2174/1570162x113116660059 (2013).

46 Pollara, J. et al. HIV-1 vaccine-induced C1 and V2 Env-specific antibodies synergize for increased antiviral activities. Journal of virology 88, 7715–7726, doi:10.1128/jvi.00156-14 (2014).

47 Ackerman, M. E. et al. Route of immunization defines multiple mechanisms of vaccine-mediated protection against SIV. Nature medicine 24, 1590–1598, doi:10.1038/s41591-018-0161-0 (2018).

48 Pardi, N. et al. Characterization of HIV-1 Nucleoside-Modified mRNA Vaccines in Rabbits and Rhesus Macaques. Mol Ther Nucleic Acids 15, 36–47, doi:10.1016/j.omtn.2019.03.003 (2019).

49 Liao, H. X. et al. Co-evolution of a broadly neutralizing HIV-1 antibody and founder virus. Nature 496, 469–476, doi:10.1038/nature12053 (2013).

50 Williams, W. B. et al. Initiation of HIV neutralizing B cell lineages with sequential envelope immunizations. Nature communications 8, 1732, doi:10.1038/s41467-017-01336-3 (2017).

51 de Taeye, S. W. et al. Immunogenicity of Stabilized HIV-1 Envelope Trimers with Reduced Exposure of Non-neutralizing Epitopes. Cell 163, 1702–1715, doi:10.1016/j.cell.2015.11.056 (2015).

52 Stamatatos, L., Pancera, M. & McGuire, A. T. Germline-targeting immunogens. Immunol Rev 275, 203–216, doi:10.1111/imr.12483 (2017).

53 LaBranche, C. C. et al. Neutralization-guided design of HIV-1 envelope trimers with high affinity for the unmutated common ancestor of CH235 lineage CD4bs broadly neutralizing antibodies. PLoS Pathog 15, e1008026, doi:10.1371/journal.ppat.1008026 (2019).

54 Pacheco, B. et al. Residues in the gp41 Ectodomain Regulate HIV-1 Envelope Glycoprotein Conformational Transitions Induced by gp120-Directed Inhibitors. J Virol 91, doi:10.1128/JVI.02219-16 (2017).

55 Zhang, P. et al. Interdomain Stabilization Impairs CD4 Binding and Improves Immunogenicity of the HIV-1 Envelope Trimer. Cell Host Microbe 23, 832–844 e836, doi:10.1016/j.chom.2018.05.002 (2018).

56 Bradley, T. et al. Amino Acid Changes in the HIV-1 gp41 Membrane Proximal Region Control Virus Neutralization Sensitivity. EBioMedicine 12, 196–207, doi:10.1016/j.ebiom.2016.08.045 (2016).

57 Gao, F. et al. Cooperation of B cell lineages in induction of HIV-1-broadly neutralizing antibodies. Cell 158, 481–491, doi:10.1016/j.cell.2014.06.022 (2014).

58 Duthie, M. S. et al. A phase 1 antigen dose escalation trial to evaluate safety, tolerability and immunogenicity of the leprosy vaccine candidate LepVax (LEP-F1 + GLA-SE) in healthy adults. Vaccine 38, 1700–1707, doi:10.1016/j.vaccine.2019.12.050 (2020).

59 Inaba, K., Metlay, J. P., Crowley, M. T. & Steinman, R. M. Dendritic cells pulsed with protein antigens in vitro can prime antigen-specific, MHC-restricted T cells in situ. The Journal of experimental medicine 172, 631–640, doi:10.1084/jem.172.2.631 (1990).

60 Inaba, K., Metlay, J. P., Crowley, M. T., Witmer-Pack, M. & Steinman, R. M. Dendritic cells as antigen presenting cells in vivo. International reviews of immunology 6, 197–206, doi:10.3109/08830189009056630 (1990).

61 Kranz, L. M. et al. Systemic RNA delivery to dendritic cells exploits antiviral defence for cancer immunotherapy. Nature 534, 396–401, doi:10.1038/nature18300 (2016).

62 Kwong, P. D. & Mascola, J. R. HIV-1 Vaccines Based on Antibody Identification, B Cell Ontogeny, and Epitope Structure. Immunity 48, 855–871, doi:10.1016/j.immuni.2018.04.029 (2018).

63 Locci, M. et al. Human circulating PD-1+CXCR3-CXCR5+ memory Tfh cells are highly functional and correlate with broadly neutralizing HIV antibody responses. Immunity 39, 758–769, doi:10.1016/j.immuni.2013.08.031 (2013).

64 Robb, M. L. et al. Risk behaviour and time as covariates for efficacy of the HIV vaccine regimen ALVAC-HIV (vCP1521) and AIDSVAX B/E: a post-hoc analysis of the Thai phase 3 efficacy trial RV 144. The Lancet. Infectious diseases 12, 531–537, doi:10.1016/s1473-3099(12)70088-9 (2012).

65 Barouch, D. H. et al. Evaluation of a mosaic HIV-1 vaccine in a multicentre, randomised, double-blind, placebo-controlled, phase 1/2a clinical trial (APPROACH) and in rhesus monkeys (NHP 13-19). Lancet (London, England) 392, 232–243, doi:10.1016/s0140-6736(18)31364-3 (2018).

66 Wang, J., Peng, Y., Xu, H., Cui, Z. & Williams, R. O., 3rd. The COVID-19 Vaccine Race: Challenges and Opportunities in Vaccine Formulation. AAPS PharmSciTech 21, 225, doi:10.1208/s12249-020-01744-7 (2020).

67 Aldon, Y. et al. Rational Design of DNA-Expressed Stabilized Native-Like HIV-1 Envelope Trimers. Cell Rep 24, 3324–3338 e3325, doi:10.1016/j.celrep.2018.08.051 (2018).

68 Saunders, K. O. et al. Vaccine Induction of Heterologous Tier 2 HIV-1 Neutralizing Antibodies in Animal Models. Cell Rep 21, 3681–3690, doi:10.1016/j.celrep.2017.12.028 (2017).

69 Joyce, M. G. et al. Soluble Prefusion Closed DS-SOSIP.664-Env Trimers of Diverse HIV-1 Strains. Cell Rep 21, 2992–3002, doi:10.1016/j.celrep.2017.11.016 (2017).

70 Zhou, T. et al. Quantification of the Impact of the HIV-1-Glycan Shield on Antibody Elicitation. Cell Rep 19, 719–732, doi:10.1016/j.celrep.2017.04.013 (2017).

71 Henderson, R. et al. Disruption of the HIV-1 Envelope allosteric network blocks CD4-induced rearrangements. Nature communications 11, 520, doi:10.1038/s41467-019-14196-w (2020).

72 Pardi, N., Muramatsu, H., Weissman, D. & Karikó, K. In vitro transcription of long RNA containing modified nucleosides. Methods in molecular biology (Clifton, N.J.) 969, 29–42, doi:10.1007/978-1-62703-260-5_2 (2013).

73 Thess, A. et al. Sequence-engineered mRNA Without Chemical Nucleoside Modifications Enables an Effective Protein Therapy in Large Animals. Molecular therapy: the journal of the American Society of Gene Therapy 23, 1456–1464, doi:10.1038/mt.2015.103 (2015).

74 Weissman, D., Pardi, N., Muramatsu, H. & Karikó, K. HPLC purification of in vitro transcribed long RNA. Methods in molecular biology (Clifton, N.J.) 969, 43–54, doi:10.1007/978-1-62703-260-5_3 (2013).

75 Maier, M. A. et al. Biodegradable lipids enabling rapidly eliminated lipid nanoparticles for systemic delivery of RNAi therapeutics. Molecular therapy: the journal of the American Society of Gene Therapy 21, 1570–1578, doi:10.1038/mt.2013.124 (2013).

76 Jayaraman, M. et al. Maximizing the potency of siRNA lipid nanoparticles for hepatic gene silencing in vivo. Angewandte Chemie (International ed. in English) 51, 8529–8533, doi:10.1002/anie.201203263 (2012).

77 Laczko, D. et al. A Single Immunization with Nucleoside-Modified mRNA Vaccines Elicits Strong Cellular and Humoral Immune Responses against SARS-CoV-2 in Mice. Immunity 53, 724–732, doi:10.1016/j.immuni.2020.07.019. (2020).

78 Hulot, S. L. et al. Comparison of Immunogenicity in Rhesus Macaques of Transmitted-Founder, HIV-1 Group M Consensus, and Trivalent Mosaic Envelope Vaccines Formulated as a DNA Prime, NYVAC, and Envelope Protein Boost. J Virol 89, 6462–6480, doi:10.1128/JVI.00383-15 (2015).

79 Saunders, K. O. et al. Vaccine Elicitation of High Mannose-Dependent Neutralizing Antibodies against the V3-Glycan Broadly Neutralizing Epitope in Nonhuman Primates. Cell reports 18, 2175–2188, doi:10.1016/j.celrep.2017.02.003 (2017).

80 Shen, X. et al. HIV-1 gp120 and Modified Vaccinia Virus Ankara (MVA) gp140 Boost Immunogens Increase Immunogenicity of a DNA/MVA HIV-1 Vaccine. Journal of virology 91, doi:10.1128/JVI.01077-17 (2017).

81 Pollara, J. et al. High-throughput quantitative analysis of HIV-1 and SIV-specific ADCC-mediating antibody responses. Cytometry. Part A: the journal of the International Society for Analytical Cytology 79, 603–612, doi:10.1002/cyto.a.21084 (2011).

82 Sambor, A. et al. Establishment and maintenance of a PBMC repository for functional cellular studies in support of clinical vaccine trials. Journal of immunological methods 409, 107–116, doi:10.1016/j.jim.2014.04.005 (2014).

83 Trkola, A., Matthews, J., Gordon, C., Ketas, T. & Moore, J. P. A cell line-based neutralization assay for primary human immunodeficiency virus type 1 isolates that use either the CCR5 or the CXCR4 coreceptor. Journal of virology 73, 8966–8974 (1999).

84 Ferrari, G. et al. An HIV-1 gp120 envelope human monoclonal antibody that recognizes a C1 conformational epitope mediates potent antibody-dependent cellular cytotoxicity (ADCC) activity and defines a common ADCC epitope in human HIV-1 serum. Journal of virology 85, 7029–7036, doi:10.1128/jvi.00171-11 (2011).

85 Tay, M. Z. et al. Antibody-Mediated Internalization of Infectious HIV-1 Virions Differs among Antibody Isotypes and Subclasses. PLoS pathogens 12, e1005817, doi:10.1371/journal.ppat.1005817 (2016).

86 Ackerman, M. E. et al. A robust, high-throughput assay to determine the phagocytic activity of clinical antibody samples. Journal of immunological methods 366, 8–19, doi:10.1016/j.jim.2010.12.016 (2011).

87 O’Doherty, U., Swiggard, W. J. & Malim, M. H. Human immunodeficiency virus type 1 spinoculation enhances infection through virus binding. J Virol 74, 10074–10080, doi:10.1128/jvi.74.21.10074-10080.2000 (2000).

88 Montefiori, D. C. Measuring HIV neutralization in a luciferase reporter gene assay. Methods in molecular biology (Clifton, N.J.) 485, 395–405, doi:10.1007/978-1-59745-170-3_26 (2009).

89 Bonsignori, M. et al. Analysis of a clonal lineage of HIV-1 envelope V2/V3 conformational epitope-specific broadly neutralizing antibodies and their inferred unmutated common ancestors. Journal of virology 85, 9998–10009, doi:10.1128/JVI.05045-11 (2011).

