## Supplemental Tables and Figures for "Lipid nanoparticle encapsulated nucleoside-modified mRNA vaccines elicit polyfunctional HIV-1 antibodies comparable to proteins in nonhuman primates"

### Supplementary Figure 1

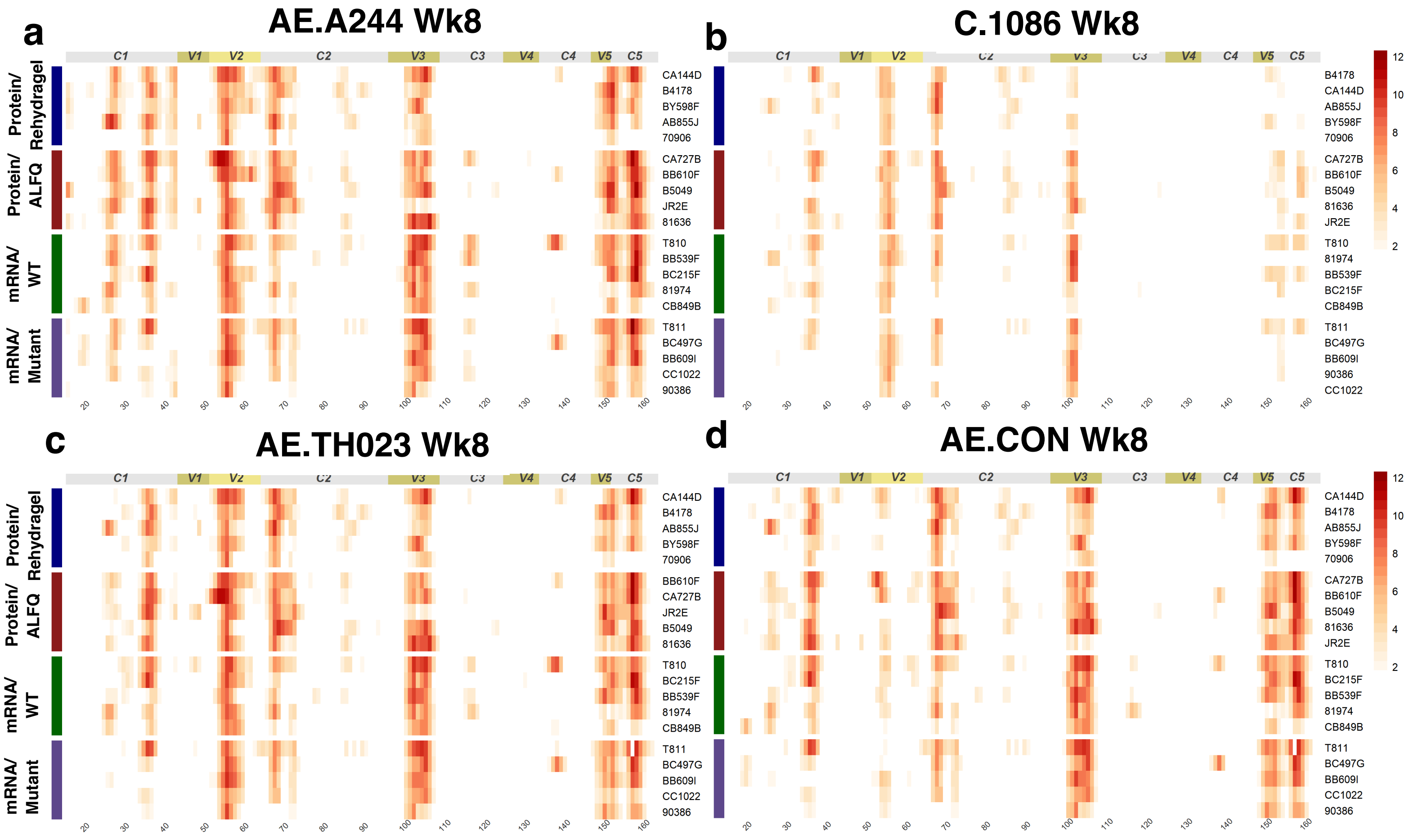


**Supplementary Figure 1. mRNA-LNP and adjuvanted recombinant protein vaccination elicits plasma IgG with cross clade specificities. a-d**. Heatmap of week 8 plasma IgG binding to peptides spanning the gp120 of **a.** AE.A244, **b.** C.1086, **c.** AE.TH023, and **d.** AE.CON. Each row shows the binding for an individual macaque. The macaque names are shown at the right side of the graph. Segments of HIV-1 gp120 are indicated across the top of the heatmap.

### Supplementary Figure 2

**a**

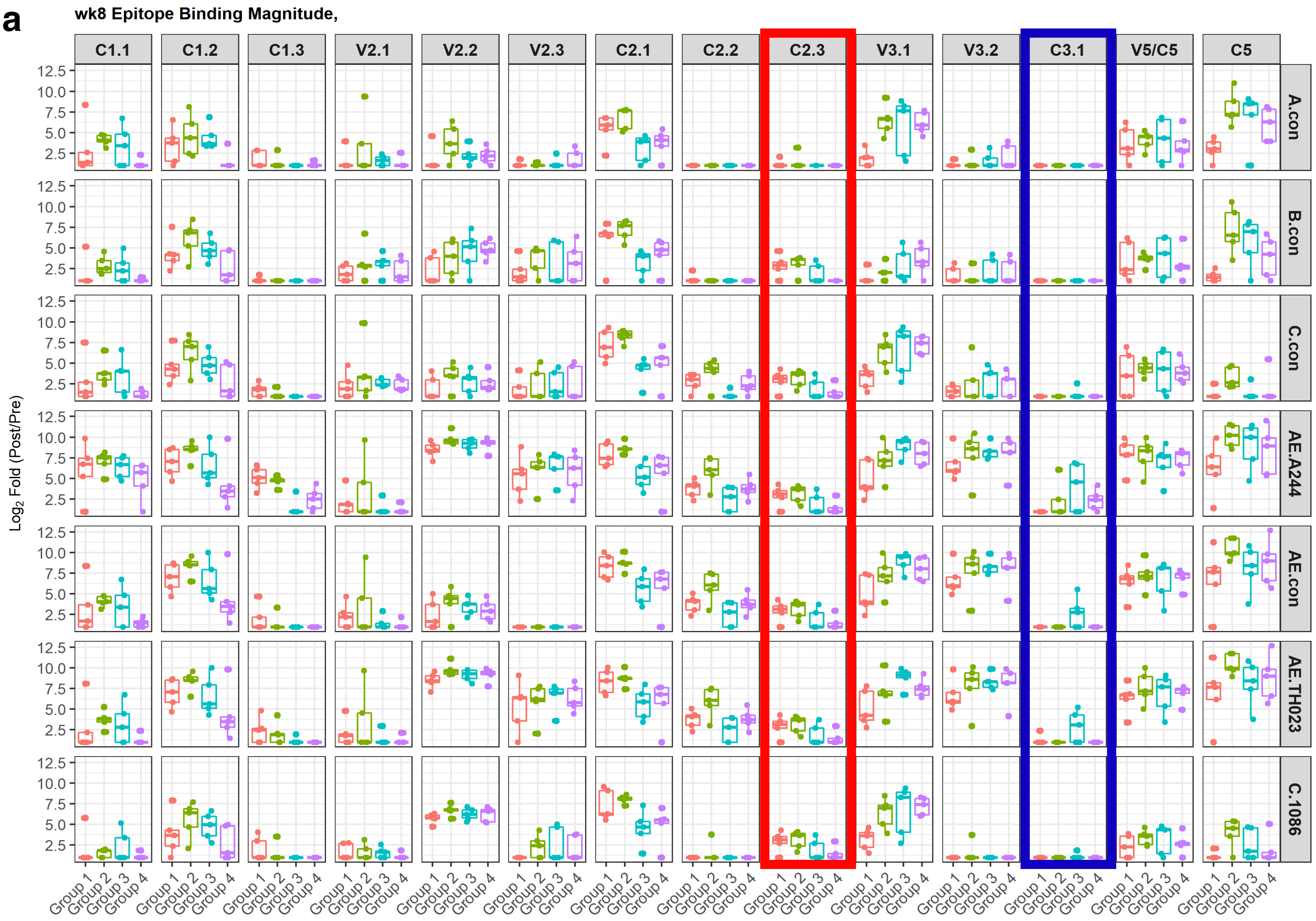


**b**

| Epitope | Start Pep# | End Pep# | HXB2_Nbr |
|---|---|---|---|
| **C1.1** | 24 | 26 | aa71-77 |
| **C1.2** | 34 | 36 | aa101-107 |
| **C1.3** | 39 | 41 | aa116-122 |
| **V2.1** | 50 | 51 | aa154-157 |
| **V2.2** | 53 | 54 | aa163-166 |
| **V2.3** | 56 | 57 | aa172-175 |
| **C2.1** | 66 | 67 | aa202-205 |
| **C2.2** | 70 | 71 | aa214-217 |
| **C2.3** | 84 | 86 | aa256-262 |
| **V3.1** | 100 | 101 | aa304-307 |
| **V3.2** | 103 | 105 | aa315-321 |
| **C3.1** | 115 | 117 | aa350-358 |
| **V5/C5** | 149 | 153 | aa463-475 |
| **C5** | 156 | 158 | aa484-490 |

**Supplementary Figure 2. Clade-specific peptide binding antibody responses post mRNA-LNP and adjuvanted recombinant protein vaccination. a**. Box-and-whisker plots of antibody binding magnitude for various Envs from different clades**.** The virus name is shown on the right. Env region is shown on the top of each column. Rectangles indicate differences in C2 (red) and C3 (blue) binding antibodies between mRNA and protein immunization. Env regions encompassed by the peptides are listed at the top of the column. **b.** Peptide locations within reference HXB2 envelope amino acid sequence.

### Supplementary Figure 3

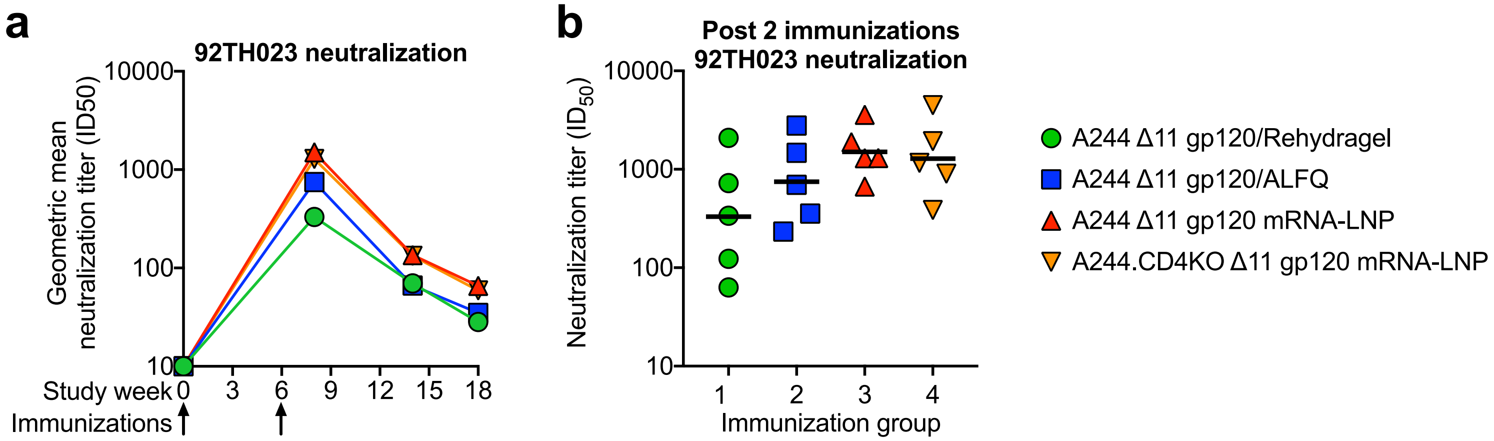


**Supplementary Figure 3. Adjuvanted recombinant gp120 and mRNA-LNP-encoded gp120 immunizations elicit comparable HIV-1 serum HIV-1.92TH023 neutralizing antibodies. a.** Serum neutralization of HIV-1.92TH023.6 infection of TZM-bl cells over the course of vaccination. The symbol represents group geometric mean. **b.** Comparison of neutralization ID50 titers two weeks post the second immunization. Neutralization titer is shown as ID50 reciprocal serum dilution. Horizontal bar represents group geometric mean.

### Supplementary Table 1

**Supplementary Table 1. Rhesus serum HIV-1 neutralization ID50 titers after 6-valent sequential SOSIP gp140 mRNA-LNP immunization.**

| | | Monkey | | | |
|---|---|---|---|---|---|
| **Week** | **Virus** | **150252** | **150794** | **150796** | **150798** |
| 18 | CH0505.w4.3 | 482 | 146 | 210 | 330 |
| 18 | CH0505 TF | 10 | 10 | 10 | 10 |
| 18 | CH0505s/293S/GnTI- | 10 | 10 | 10 | 10 |
| 18 | CH0505TF.gly4 | 10 | 10 | 10 | 10 |
| 18 | CH0505s.G458Y.4 | 10 | 10 | 10 | 10 |
| 18 | CH0505s.G458Y.4/293S/GnTI- | 10 | 10 | 10 | 10 |
| 32 | CH0505s.G458Y.N279K.2/293S/GnTI- | 10 | 48 | 32 | 10 |
| 32 | CH0505TF.N279K/293S/GnTI- (M5) | 10 | 10 | 10 | 10 |
| 18 | TRO.11 | 10 | 10 | 10 | 10 |
| 18 | SVA-MLV | 10 | 10 | 10 | 10 |

Week 18 and 32 are two and 16 weeks after the fifth and final immunization respectively. SVA-MLV, murine leukemia virus.

### Supplementary Table 2

**Supplementary Table 2. Rhesus serum HIV-1 neutralization ID50 titers 6-valent sequential gp160 mRNA-LNP immunization.**

| | | Monkey | | | |
|---|---|---|---|---|---|
| **Week** | **Virus** | **150250** | **150793** | **150795** | **6858** |
| 18 | CH0505.w4.3 | 1164 | 3111 | 4169 | 2495 |
| 18 | CH0505 TF | 10 | 10 | 10 | 10 |
| 18 | 001428-2.42 | 10 | 10 | 10 | 10 |
| 18 | 426c.N276D.N460D.N463D/293S/GnTI- | 10 | 43 | 10 | 10 |
| 18 | 426c.N460D.N463D/293S/GnTI- | 10 | 10 | 10 | 10 |
| 18 | CH0505.w24.e12 (CH505.30.12) | 10 | 10 | 10 | 10 |
| 18 | CH0505.w24.e20 (CH505.30.20) | 10 | 10 | 10 | 10 |
| 18 | CH0505TF.M11 (CH505.M11) | 10 | 10 | 10 | 10 |
| 18 | CH0505TF.M5 (CH0505.M5) | 10 | 10 | 10 | 10 |
| 18 | CH0505TF.gly3.276 | 10 | 10 | NT | 10 |
| 18 | HIV CH0505.C.e.B18 (CH505.136.B18) | 10 | 10 | 10 | 10 |
| 18 | HIV CH0505TF.gly4 | 10 | 10 | 10 | 10 |
| 18 | SVA-MLV | 10 | 10 | 10 | 10 |

Week 18 is two weeks after the fifth and final immunization. NT, not tested due to sample availability. SVA-MLV, murine leukemia virus.

### Supplementary Table 3

**Supplementary Table 3. Rhesus serum HIV-1 neutralization ID50 titers after 6-valent sequential SOSIP gp140 protein/GLA-SE immunization.**

| | | Monkey | | | |
|---|---|---|---|---|---|
| **Week** | **Virus** | **150251** | **6857** | **T244** | **T245** |
| 18 | CH0505.w4.3 | 450 | 122 | 783 | 200 |
| 18 | CH0505 TF | 10 | 10 | 10 | 10 |
| 32 | CH0505TF.N279K/293S/GnTI- (M5) | 10 | 10 | 10 | 10 |
| 18 | CH0505TF.gly4 | 10 | 62 | 10 | 10 |
| 18 | CH0505s.G458Y.4 | 10 | 10 | 10 | 10 |
| 18 | CH0505s.G458Y.4/293S/GnTI- | 37 | 10 | 10 | 10 |
| 32 | CH0505s.G458Y.N279K.2/293S/GnTI- | NT | NT | 10 | 24 |
| 18 | CH0505s/293S/GnTI- | 46 | 10 | 32 | 10 |
| 18 | TRO.11 | 10 | 10 | 10 | 10 |
| 18 | SVA-MLV | 10 | 10 | 10 | 10 |

Week 18 and 32 are two and 16 weeks after the fifth and final immunization respectively. NT, not tested due to sample availability. SVA-MLV, murine leukemia virus.
